# A basement membrane discovery pipeline uncovers network complexity, new regulators, and human disease associations

**DOI:** 10.1101/2021.10.25.465762

**Authors:** Ranjay Jayadev, Mychel RPT Morais, Jamie M Ellingford, Sandhya Srinivasan, Richard W Naylor, Craig Lawless, Anna S Li, Jack F Ingham, Eric Hastie, Qiuyi Chi, Maryline Fresquet, Nikki-Maria Koudis, Huw B Thomas, Raymond T O’Keefe, Emily Williams, Antony Adamson, Helen M Stuart, Siddharth Banka, Damian Smedley, Genomics England Research Consortium, David R Sherwood, Rachel Lennon

## Abstract

Basement membranes (BMs) are ubiquitous extracellular matrices whose composition remains elusive, limiting our understanding of BM regulation and function. By developing a bioinformatic and *in vivo* discovery pipeline, we define a network of 222 human proteins localized to BMs. Network analysis and screening in *C. elegans* and zebrafish identified new BM regulators, including ADAMTS, ROBO, and TGFβ. More than 100 BM-network genes associate with human phenotypes and by screening 63,039 genomes from families with rare disorders, we discovered loss-of-function variants in *LAMA5*, *MPZL2*, and *MATN2,* and show they regulate BM composition and function. This cross-disciplinary study establishes the immense complexity and role of BMs in human health.

---

Basement membranes (BMs) are the most ancient animal extracellular matrix (ECM) and form sheet-like structures that underlie epithelia and surround most tissues^1, 2^. Two independent planar networks of laminin and type IV collagen molecules associate with cell surface interactors (CSIs) and provide the scaffolding structure that builds BMs along tissues^2–5^. The glycoprotein nidogen and the heparan sulfate proteoglycan perlecan bridge the laminin and collagen IV scaffolds^6^. BMs also harbor matricellular proteins, growth factors, and proteases^1^. BMs have diverse compositions tailored to resist mechanical stress, dictate tissue shape, and create diffusion barriers^7–10^. They also provide cues that direct cell polarity, differentiation, migration, and survival^5, 11–13^. Underscoring their diverse and essential functions, variants in more than twenty BM genes underlie human diseases^14^. BM proteins are targets of autoantibodies in immune disorders^15^, and defects in BM protein expression and turnover are a key pathogenic aspect of cancer, diabetes, and fibrosis^16–19^.

The number of proteins that associate with BMs remains unclear. Gene Ontology (GO) and ECM annotations estimate between 24 to 100 BM proteins^16, 20^, while proteomic studies suggest that BMs contain significantly more than 100 proteins^20^. Challenges with protein solubility, abundance, and loss of spatial characteristics, however, have limited the use of biochemical methods in identifying BM constituents^16^. *In silico* prediction has identified more than 1,000 putative matrisome (ECM) proteins in vertebrates and over 700 in *C. elegans*^21–23^. As there are many different ECMs, it is unknown whether many of these components bind to, function within, or regulate BMs. Validation of a candidate BM protein requires its microscopic localization to a BM *in vivo*^20^. Many matrix proteins have been detected within vertebrate BMs by immunolocalization^24–27^ and *C. elegans* studies have characterized more than 30 fluorophore-tagged endogenous BM proteins^4, 28^. However, without a focused effort to curate these proteins and the development of new strategies to identify BM proteins, our understanding of BMs in normal and disease states remains limited.

In this study we develop a bioinformatic and *in vivo* localization pipeline and define a comprehensive network of 222 human genes encoding BM matrix proteins and BM cell surface interactors (collectively referred to as BM zone genes), many of which show tissue-specific expression and BM localization in humans and mice. Using domain network analysis, we identify key regulatory hub proteins, including perlecan and papilin, which maintain BM structure and promote BM turnover. Through knockdown studies in *C. elegans* and zebrafish, we reveal that Robo receptors and TGFβ signaling limit BM type IV collagen levels. We further identify human gene-phenotype associations for 112 BM-network genes. By screening the rare disease cohort in the Genomics England 100,000 Genomes Project (100KGP)^29^, we uncover new disease associations for rare loss-of-function variants in *MATN2, MPZL2*, and *LAMA5* and demonstrate these variants are linked to defective BM composition, structure, and function. Collectively, this cross-disciplinary approach provides new insights into BM assembly, regulation, and disease, and establishes the vast complexity of BMs and their critical importance in human health.

## Results

### A comprehensive BM gene network across species

The number of BM-associated proteins has not been rigorously determined. To establish a comprehensive network of BM zone genes, we first identified 103 human “basement membrane”-annotated genes from the Gene Ontology (GO) Resource^30, 31^. We then curated data across species for known BM components and predicted new candidates through gene expression, protein interaction, and domain enrichment analyses (**Methods**, **Supplementary notes**, and **Supplementary tables 1-5**), which yielded 160 additional BM zone candidates (**Supplementary table 6**). Next, we confirmed BM zone localization for 174 out of 263 candidates based on protein immunolocalization studies and from 36 fluorescent tagged orthologs previously localized to *C. elegans* BM (**Fig. 1a-b**; **Supplementary Fig. 1a** and **Supplementary tables 5-7**). To further verify BM association, 25 *C. elegans* BM gene orthologs were endogenously tagged with mNeonGreen (mNG) (**Supplementary notes** and **Supplementary Fig. 2a**). We examined all post-embryonic BMs and highlight the pharynx (feeding organ) and gonad BMs (**Fig. 1c**, EMB-9::mRuby2 (COL4A1)). Among proteins with only predicted BM localization, the UNC-5 (UNC5) receptor polarized to both pharyngeal and gonadal BMs, and the membrane-associated C16E9.1 (MATN1) polarized to the pharyngeal BM but was not detected in the gonad (**Fig. 1c** and **Supplementary table 5**). For proteins with limited BM immunolocalization evidence, TEST-1 (SPOCK), CPI-1 (CST3), CPI-2 (CST3), and SAX-3 (ROBO) localized to both BMs (**Fig. 1c** and **Supplementary table 5**), while ADM-4 (ADAM17), C48E7.6 (CSPG4), and F26E4.7 (POSTN/TGFBI) localized to the pharyngeal BM (**Supplementary Fig. 2b**). NAS-39 (TLL1), EVA-1 (EVA1), and FBN-1 (FBN) were not detected in BMs but were visible in other ECMs (**Supplementary Fig. 2c** and **Supplementary table 5**). Fluorescent tagging localized 14 additional candidates, and together these verification strategies confirmed BM zone localization for 188 proteins (**Fig. 1**). Localization for 34 of 75 remaining candidates was predicted based on protein interaction with verified BM zone proteins and BM-cleaving protease activity, and 41 proteins without verified or predicted BM localization were not considered for further analyses (**Fig. 1a**, **Supplementary Fig 1a,** and **Supplementary tables 6-7**). This pipeline defined an expanded network of 160 mammalian BM matrix proteins and 62 CSIs that localize to the BM zone, with orthologs in *C. elegans*, zebrafish, and *Drosophila* (**Fig. 1b**, **Supplementary Fig. 1b and 3**, and **Supplementary Table 7**). To curate the BM zone network, we established the ***<u>b</u>****asement **<u>m</u>**embraneBASE* database (bmBASE; https://bmbase.manchester.ac.uk).

**Figure 1.**
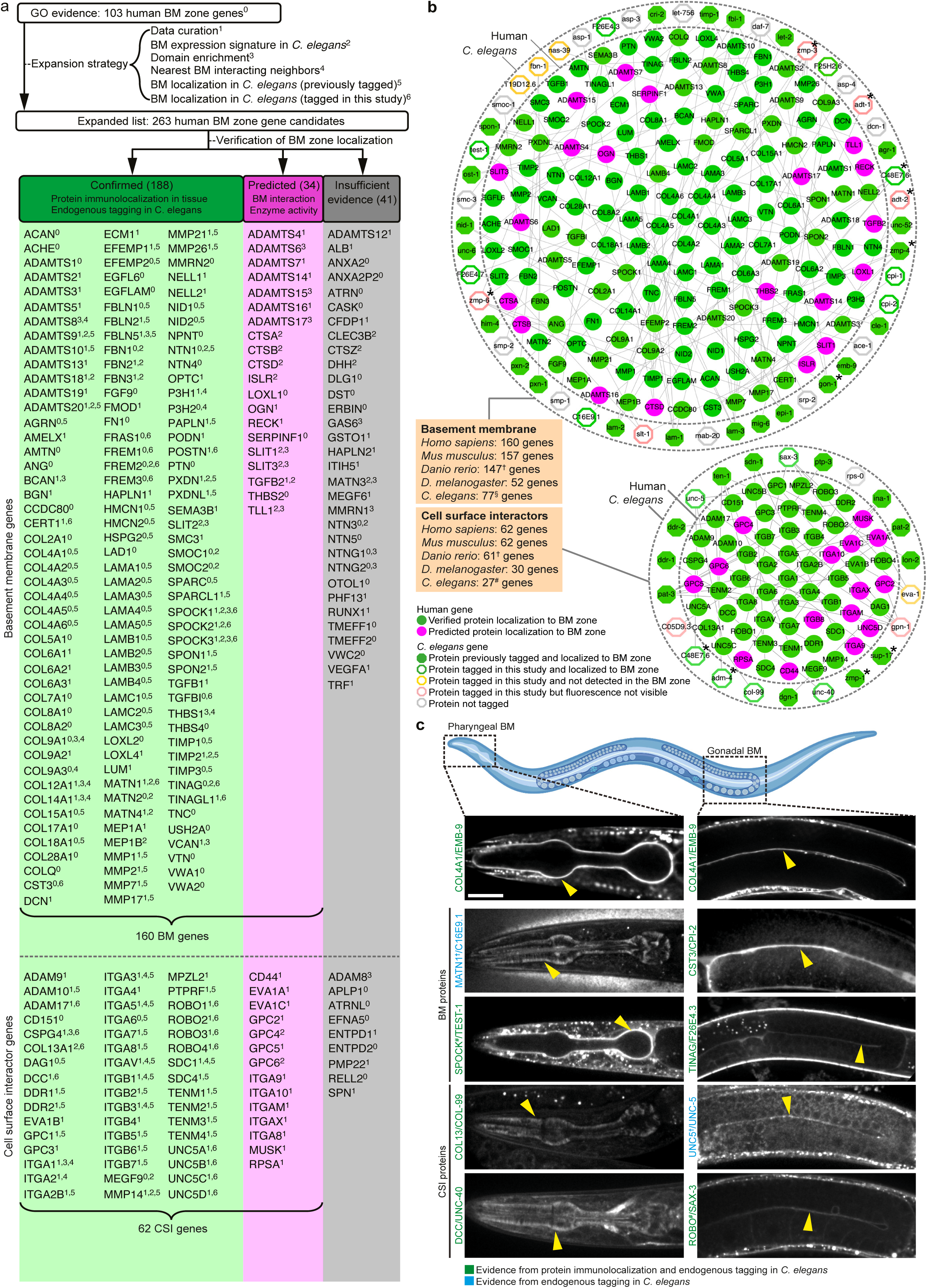
A comprehensive and conserved network of basement membrane zone genes. **a**, Strategy for identifying putative basement membrane (BM) matrix protein and cell surface interactor (CSI) genes in humans and other species. Protein localization to BM zone was either (i) confirmed through vertebrate tissue immunolocalization and/or fluorescent tagging in *C. elegans* (green column), or (ii) predicted based on BM protein interaction or BM-cleaving protease activity (magenta column). Genes with insufficient evidence are in the grey column. **b**, The integrated BM zone network of 222 human genes (circles, 160 BM matrix and 62 CSI components) with corresponding *C. elegans* orthologs (octagons). Lines link ortholog pairs. Boxes indicate number of ortholog genes in mouse, zebrafish, *Drosophila*, and *C. elegans*. *Complex ortholog families in *C. elegans* (detailed in **Supplementary fig. 3**). ^†^Duplicated zebrafish genes were counted as one; worm ^§^astacins and ^#^tetraspanins not included. Network curated at https://bmbase.manchester.ac.uk/. **c**, Top, schematic representation of the pharyngeal and gonadal BMs in *C. elegans* (<u>Biorender</u>). Middle panel, confocal mid-plane z-slices of adult animals expressing endogenously tagged COL4A1/EMB-9::mRuby2. Lower panels, selected BM zone candidates tagged with mNeonGreen in this study. Yellow arrowheads indicate BM zone localization. *Predicted human candidates that localized to the BM zone in *C. elegans*. ^#^Localization evidence present for 1/3 SPOCK and 1/4 ROBO family members in humans (**Supplementary table 7**). Scale bar represents 25 μm.

### Identification of tissue-specific BM composition signatures

To investigate BM diversity in human tissues, the relative abundance of the BM zone network within proteomic datasets for human kidney^19^, liver^16^, colon^16^, and omentum^32^ was ranked (**Supplementary table 8**). We found a common set of BM zone proteins that were abundant across all tissues, including COL4A1/2, COL6A1/A2/A3, LAMB2/C1, NID1, and HSPG2 (perlecan), and identified proteins with highly variable abundance, such as DCN and TINAG (**Fig. 2a**). Moreover, distinct patterns of protein composition and abundance within tissues were apparent (see kidney glomerulus compared to tubulointerstitium, **Fig. 2a**). Analysis of 121 human and 51 mouse tissue transcriptomes was conducted to determine whether these differences were linked to gene expression (**Supplementary tables 9 and 10**). Dynamic clustering revealed strong BM gene grouping by tissue (**Fig. 2b** and **Supplementary Fig. 4a-b**). Furthermore, there was segregation by tissue compartment (*e.g.*, cerebral cortex and cerebellum) and a distinct clustering pattern in organs with mucosal linings (*e.g.*, intestine, fallopian tubes, gallbladder, urinary bladder), suggesting that BM expression signatures associate with tissue functions (**Fig. 2b** and **Supplementary Fig. 4a**). Ranking BM genes by expression variance revealed low variance genes (expressed in most tissues) were predominantly glycoproteins and matrix regulators such as laminins, nidogens, and papilin (**Supplementary table 10** and **Fig. 2c-d**). Very high variance genes included SPARC-related genes, many collagen chains, and secreted BM-binding factors (**Fig. 2c-d**). Together, these analyses identify protein composition and gene expression patterns that contribute to BM signatures for different human tissues and reveal both common BM template components and tissue-specific BM factors.

**Figure 2.**
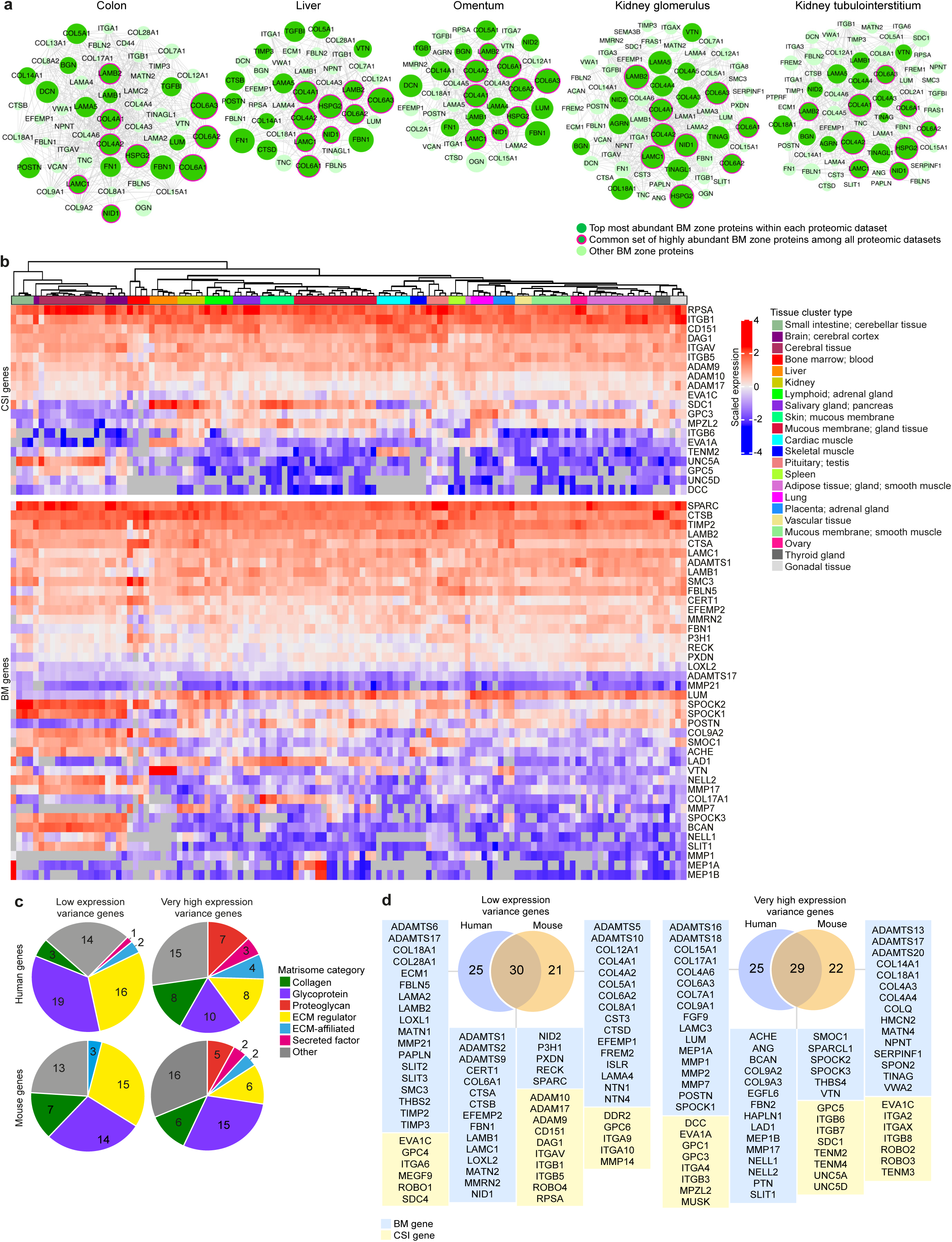
Mammalian basement membranes of different tissues have unique compositional signatures. **a**, Interactomes (based on published proteomic datasets) representing the diversity of BM composition across human tissues. Proteins are depicted as nodes sized according to their log-transformed relative tissue abundance. Lines represent protein interactions determined by STRING analysis (**Supplementary table 4**). **b**, Heatmaps derived from published transcriptomic datasets (**Supplementary table 9**) depict either very high or low expression variance of BM zone genes across multiple human tissues. **c**, Classification of human and mouse BM and CSI genes with differing expression variance according to matrisome category. **d**, Overlap between human BM zone genes and mouse orthologs for very high or low expression variance. See **Supplementary fig. 4** and **Supplementary table 10** for extended expression variance data.

### Domain network analysis reveals BM hub proteins

How BM zone proteins are organized and interact is not well understood. To investigate these relationships, we mapped protein domains enriched in our BM zone network and generated human and *C. elegans* domain-based networks (**Fig. 3a-b** and **Supplementary Fig. 5a-b**). We found distinct protein sub-networks based on domain commonality (*e.g.,* integrin subunits, ADAMTS proteases, and collagen IV chains) and a broader module including laminins, other glycoproteins, and proteoglycans (**Fig. 3b**). The same clustering pattern was present in *C. elegans*, indicating shared modular assembly and domain connectivity (**Supplementary Fig. 5b**).

**Figure 3.**
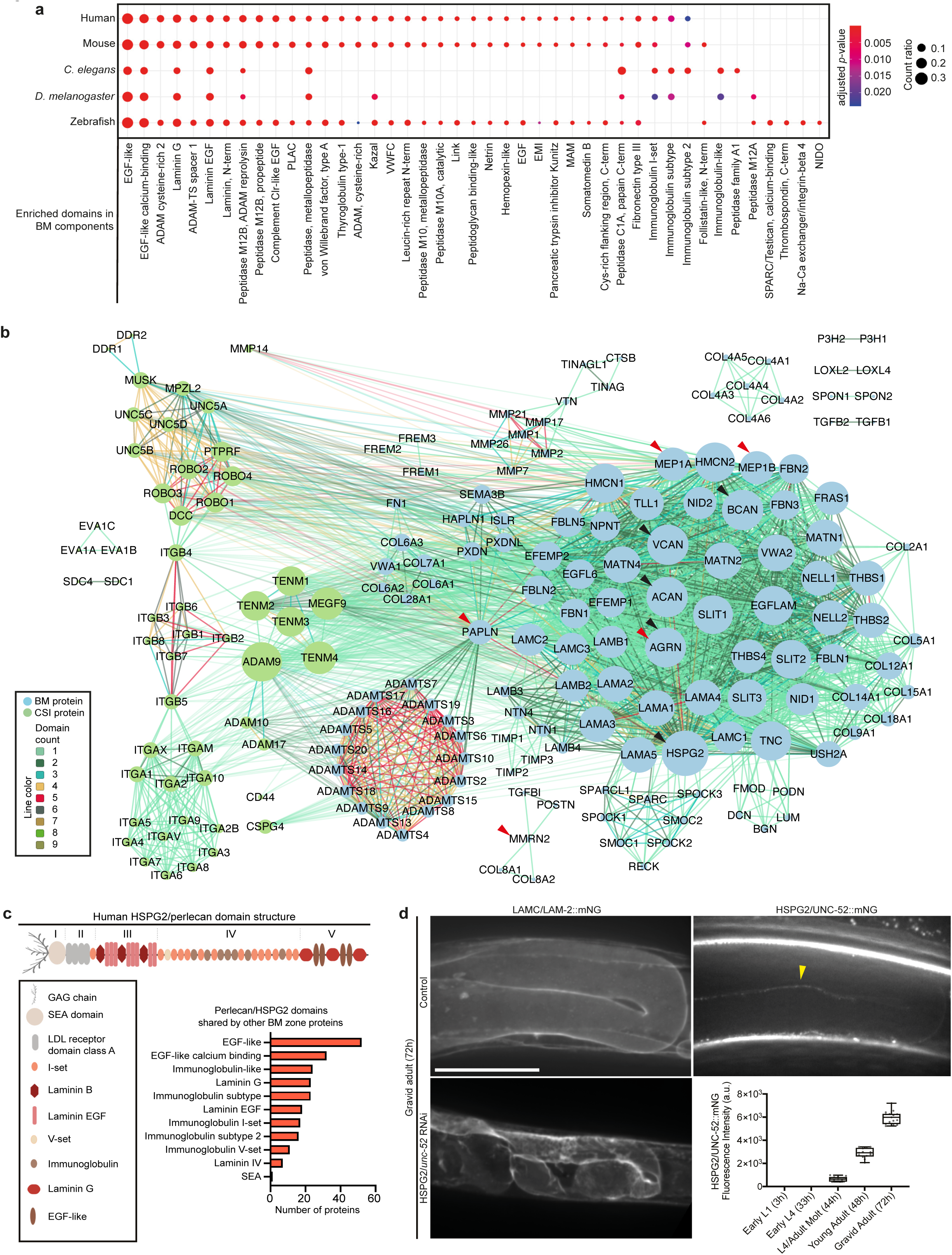
Domain network analysis reveals hub proteins in the basement membrane zone. **a**, Dot plot depicts conserved and enriched Interpro protein domains in BM zone components. Dots are sized according to number of domain occurrences within BM genes (count ratio, **Supplementary table 3**). **b**, A domain-based interactome for human BM zone genes. Nodes represent BM and CSI proteins and are sized according to network degree score. Lines connecting nodes are color-coded to indicate the number of shared domains. Black and red arrowheads highlight hub proteins with the highest degree and betweenness centrality scores respectively (**Methods** and **Supplementary table 11**). **c**, Domain structure of the hub protein HSPG2/perlecan. Bar chart shows the number of BM zone proteins sharing specific domains with HSPG2. **d**, Left, confocal sum projections of *C. elegans* gonadal BM LAMC/LAM-2::mNG in control vs. HSPG2*/unc-52* RNAi-treated 72-h adult animals (n = 10 animals examined each). Top right, middle-plane confocal z-slice of HSPG2/UNC-52::mNG in an adult animal, yellow arrowhead indicates fluorescence signal in the gonadal BM; bottom right, quantification of gonadal BM UNC-52::mNG levels throughout post-embryonic development below (n ≥ 13 for each developmental stage; fluorescence was not detected between early L1 and early L4 stages). Scale bar represents 25 μm. For boxplots, edges indicate the 25^th^ and 75^th^ percentiles, the line in the box represents the median, and whiskers mark the minimum and maximum values.

Next, network connectivity metrics were calculated to identify potential regulatory hubs (**Methods**). The degree score, which indicates the number of connections between proteins based on shared domains, was determined using the Cytoscape NetworkAnalyzer tool^33^. Among BM matrix proteins, the heparan sulfate proteoglycan perlecan (HSPG2) and its *C. elegans* ortholog UNC-52 had the highest degree score, sharing specific domains with 84 BM proteins (**Supplementary table 11** and **Fig. 3c**). The precise role of perlecan in BMs is not clear as its loss causes embryonic lethality in mice and *C. elegans*^34–36^. In *Drosophila*, post-embryonic perlecan depletion results in misshapen organs, suggesting a role within BMs in shaping tissues^37–39^. We examined post-embryonic perlecan (UNC-52) in *C. elegans* and discovered its localization in the adult gonadal BM (**Fig. 3d**). Depletion of UNC-52 caused a progressive compaction of the gonad from young adulthood, correlating with the onset of perlecan localization (**Fig. 3d** and **Supplementary Fig. 5d**). Furthermore, the smooth appearance of the sheet-like BM was disrupted by fibrillar structures and aggregates of laminin, collagen IV, nidogen, and papilin (**Fig. 3d** and **Supplementary Fig. 5e**). These data indicate a shared role for perlecan in shaping tissues and reveal a function in stabilizing BM organization.

The betweenness centrality score, which ranks BM proteins that connect subnetworks, was also calculated and the proteoglycan papilin (PAPLN) and its *C. elegans* ortholog MIG-6 scored the highest (**Supplementary table 11**). PAPLN and MIG-6 connect diverse classes of BM and CSI proteins, including ADAMTS proteases, peroxidasins, various proteoglycans, DCC/UNC-40, and ROBO (**Fig. 3b** and **Supplementary Fig. 5c**). Supporting this analysis, a recent study found *C. elegans* MIG-6 promotes collagen IV turnover by restricting the BM localization of ADAMTS proteases and peroxidasin^28^. Additional BM matrix and CSI proteins with high network connectivity scores were identified, including AGRN, MEP1A, ACAN, teneurins, ROBO, and DCC (**Fig. 3b** and **Supplementary table 11**). Collectively, these domain network analyses pinpoint BM constituents that organize multiple interactions within BMs regulating key BM properties such as growth, shape, and integrity.

### Functional *in vivo* studies uncover new regulators of BM composition

Little is known about how BM composition is controlled during tissue growth. We thus performed a RNAi screen targeting 77 *C. elegans* BM zone orthologs and examined laminin and collagen IV in the gonadal BM, which expands ∼100-fold in surface area during larval development (**Supplementary table 12**). Knockdown of 19 genes affected BM composition, including previously reported integrin receptor and papilin genes^4, 28^ (**Fig. 4a** and **Supplementary table 12**). Of the newly identified regulatory genes, knockdown of *adt-2* (*ADAMTS3*) caused a 40% reduction in BM laminin and collagen IV levels, and led to misshapen gonads (**Fig. 4b** and **Supplementary table 12**). Depletion of SAX-3 (ROBO), which strongly polarizes to the gonadal BM (**Fig. 1c**), increased laminin and collagen IV levels (**Fig. 4b**). Loss of the ROBO ligand SLT-1/SLT, however, did not show a similar phenotype, suggesting a SLT-independent function for ROBO (**Fig. 4a** and **Supplementary table 12**). Finally, loss of the TGFβ ligand-encoding genes *dbl-1*, *tig-2*, *tig-3*, *daf-7*, and *unc- 129* and the two TGFβ type I receptor-encoding genes *daf-1* and *sma-6* led to elevated collagen IV levels without affecting laminin (**Fig. 4b, Supplementary Fig. 6a-b**, and **Supplementary table 12**). Consistent with a cell-autonomous role for TGFβ signaling in regulating collagen IV levels, the sole *C. elegans* TGFβ type II receptor DAF-4 polarizes towards the gonadal BM (**Supplementary Fig. 6c**).

**Figure 4.**
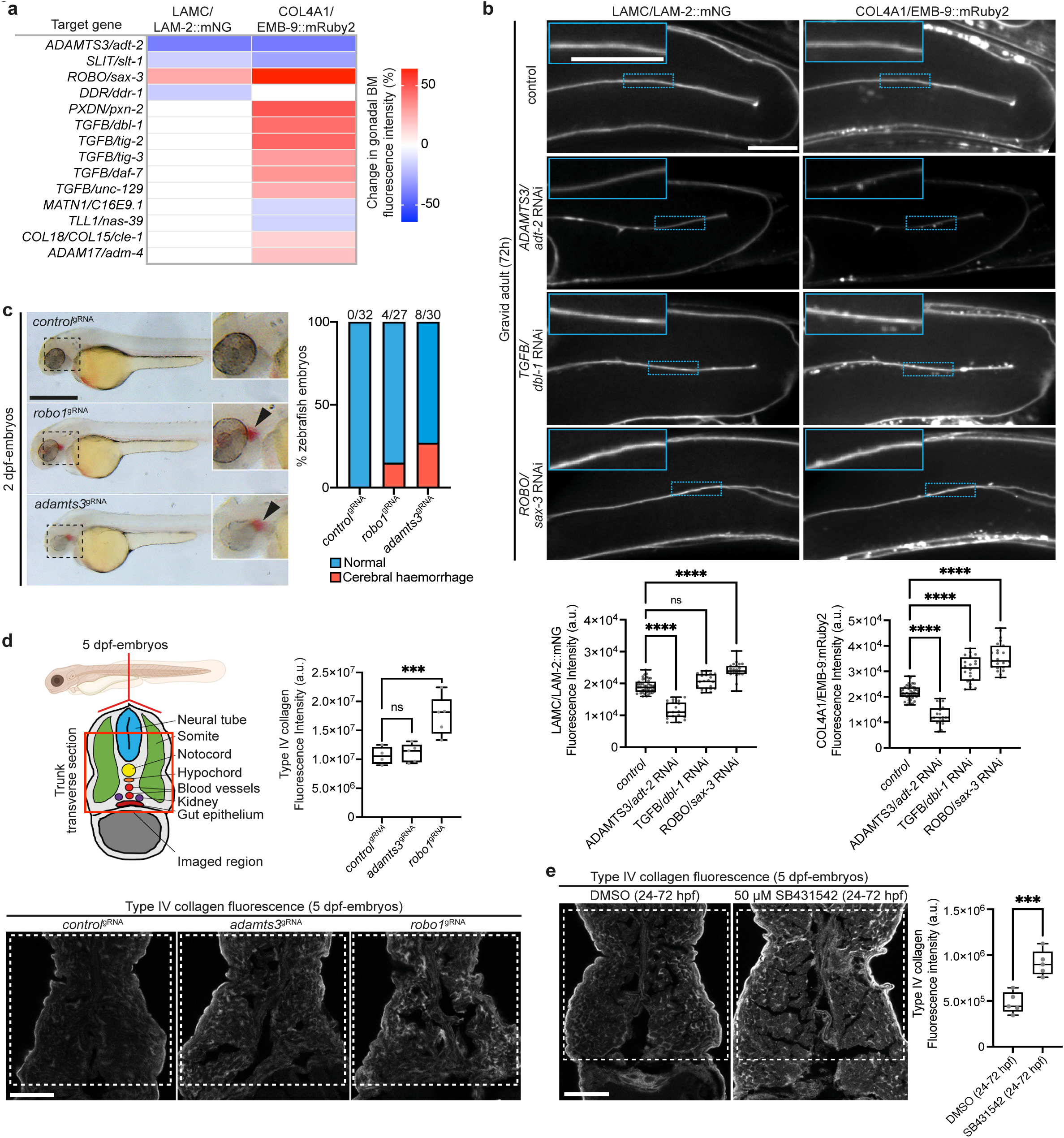
New regulators of basement membrane composition in *C. elegans* and zebrafish. **a**, Heatmap summarizing changes in LAMC/LAM-2::mNG and COL4A1/EMB- 9::mRuby2 fluorescence upon target gene knockdown in *C. elegans*. **b**, Confocal middle-plane z-slices of gonadal BM LAMC/LAM-2::mNG and COL4A1/EMB-9::mRuby2 in control and RNAi-targeted *adt-2*, *dbl-1*, and *sax-3* 72-h adult animals (boxed regions magnified in insets) with quantifications of fluorescence intensity shown below (n ≥ 20 each). \*\*\*\**p*-value < 0.0001, ns—not significant; one-way ANOVA with post-hoc Dunnett’s test. Scale bars: 25 μm. **c**, Left, brightfield images of 2-day post fertilization (dpf) zebrafish embryos injected with indicated gRNAs (boxed regions magnified in insets). Right, observed frequency of intracerebral haemorrhage (arrowheads). Scale bar represents 600 μm. **d**, Top left, schematic cross-section of a 5-dpf zebrafish embryo (<u>Biorender</u>); red box indicates imaged region. Bottom, confocal images of type IV collagen immunofluorescence in *control*^gRNA^, *adamts3*^gRNA^, and *robo1*^gRNA^-injected 5-dpf embryos; fluorescence intensity within dashed boxes is quantified on the top right (n = 5 for each treatment). \*\*\**p*-value < 0.001, ns—not significant; one-way ANOVA with post-hoc Dunnett’s test. Scale bar represents 30 μm. **e**, Collagen IV immunofluorescence in 5-dpf embryos treated with DMSO (control) or SB431542 (TGFBR1 inhibitor) on the left; quantification of fluorescence intensity within dashed boxes on the right (n = 5 each). \*\*\**p-*value < 0.001, unpaired Student’s *t* test. Scale bar represents 30 μm. For boxplots, edges indicate the 25^th^ and 75^th^ percentiles, the line in the box represents the median, and whiskers mark the minimum and maximum values.

We next investigated whether ADAMTS3, ROBO, and TGFβ have roles in regulating vertebrate BM composition. A*damts3* and *robo1* CRISPR-knockdown (crispant)^40^ zebrafish exhibited intracerebral hemorrhaging, which also occurs after *col4a1* knockdown due to vascular BM disruption, and defective fin folds, which is associated with BM dysfunction^41^ (**Fig. 4c** and **Supplementary Fig. 6d-e**). In addition, we examined kidney physiology, which depends on BMs for filtration function^9^. Using the *NL:D3* excretion assay^42^, proteinuria was observed in *adamts3* and *robo1* crispants (**Supplementary Fig. 6f**), suggesting defects in BM. Immunofluorescence studies revealed elevated collagen IV levels in *robo1* crispant zebrafish larvae, similar to ROBO loss in *C. elegans* (**Fig**. **4d**). Finally, to investigate TGFβ signaling, zebrafish larvae were treated with the TGFβ type I receptor inhibitor SB431542, which resulted in increased BM collagen IV levels, also mirroring *C. elegans* (**Fig. 4e**). Together, these *in vivo* screens support the emerging role of ADAMTS proteins in BM regulation^28, 43^ and uncover a new function for the Robo receptor and TGFβ signaling in limiting BM collagen IV levels.

### Expansion of BM disease associations

Previous studies established that germline variants in ∼30 BM genes cause genetic disorders, approximately half of which encode collagen IV and laminin chains^14^. We used our BM zone network to search for disease associations in the Human Phenotype Ontology (HPO)^44^, Genomics England PanelApp^45^, and Online Mendelian Inheritance in Man (OMIM)^46^ databases. Gene-phenotype associations with varying degrees of evidence were found for 112 BM zone genes and most were associated with autosomal recessive inheritance (**Fig. 5a-b** and **Supplementary table 13-14**). Collagen- and glycoprotein-encoding genes such as *COL2A1*, *COL4A1*, *COL7A1*, *FBN1*, *TGFBI*, and the proteoglycan *HSPG2* showed the highest number of associations with phenotypic abnormalities (top-level HPO terms, **Fig. 5c** and **Supplementary table 14**). The eye, nervous system, head and neck, skeletal, and limbs had disease phenotype associations with more than 100 BM zone genes, suggesting sensitivity of these organ systems to BM component loss (**Fig. 5c** and **Supplementary table 14**). Together, this analysis greatly expands BM gene association with human disease and reflects BMs’ diverse roles in supporting tissues.

**Figure 5.**
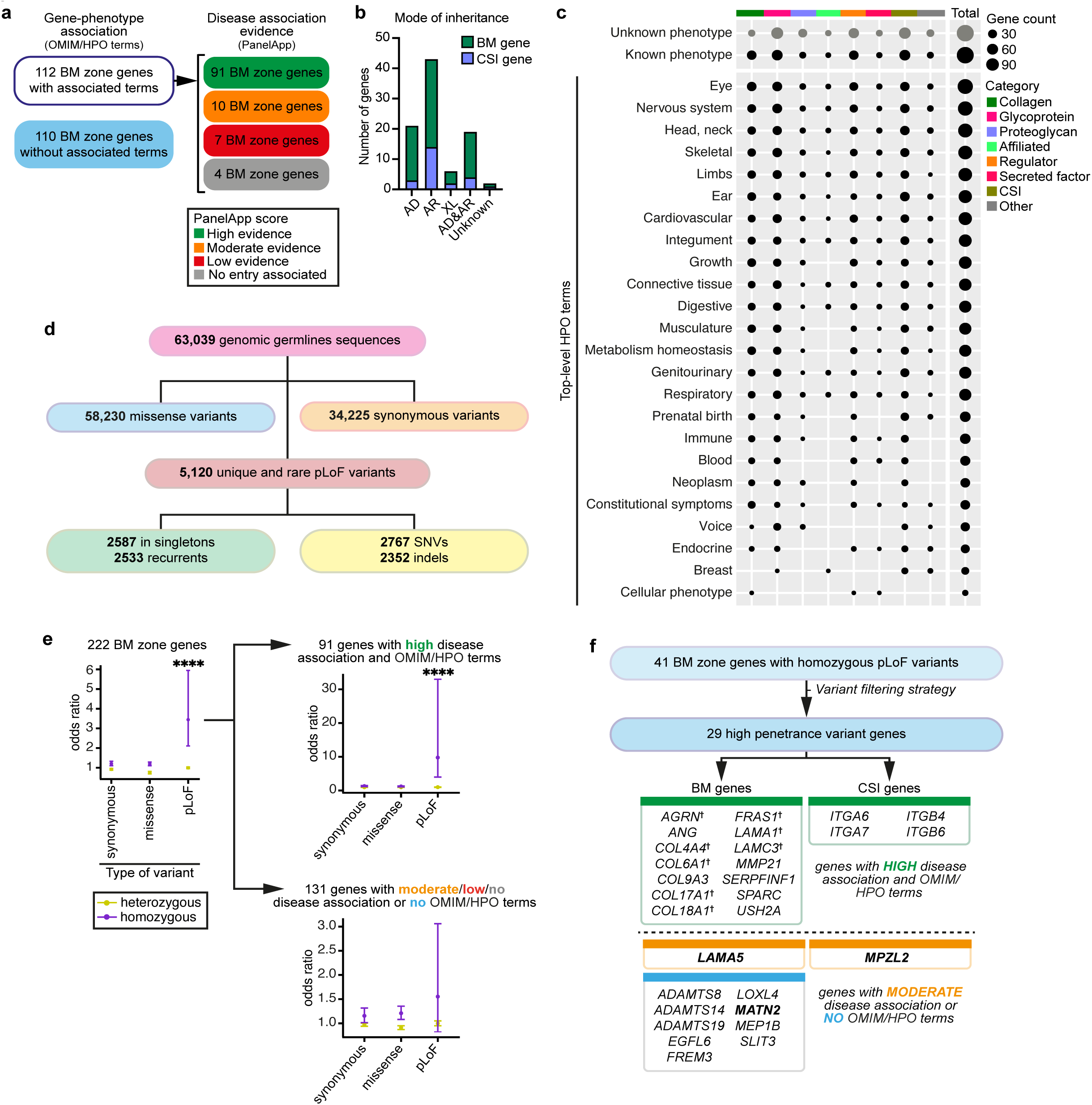
Expansion of human disease associations and identification of candidate disease-causing variants in basement membrane zone genes. **a**, Classification of BM zone genes according to phenotypes within OMIM and HPO databases (left), and level of evidence for disease causation within Genomics England PanelApp database (right); see also **Supplementary tables 13-14**). **b**, Bar chart depicting mode of inheritance for 112 BM zone genes with disease associations (AD: autosomal dominant, AR: autosomal recessive, XL: X-linked). **c**, Dot plot illustrates phenotypic abnormalities (HPO terms) associated with BM zone genes grouped by their matrisome classifications. Dot size indicates gene count in each category. **d**, Landscape of BM zone gene variants identified in the Genomics England 100KGP cohort (**Supplementary table 15**). **e**, Odds ratio (OR) plots (comparing individuals with disease to unaffected relatives) for all variants (left) or variants grouped according to disease association evidence for the respective BM zone genes (right). \*\*\*\**p*-value < 0.0001, Fisher’s Exact test. **f**, High penetrance homozygous pLoF variants were identified for 29 BM zone genes, including **^†^**components previously associated with BM-linked disease, and new disease candidates. Potential disease-causing mechanisms were investigated for genes highlighted in bold.

### Genomic studies identify candidate disease variants in BM zone genes

To determine if additional BM zone genes contribute to human diseases, 100KGP germline genomic sequences for 34,842 individuals affected with rare disease and 28,197 unaffected relatives were examined. We identified 34,225 synonymous, 58,230 missense, and 5120 predicted loss-of-function (pLoF) heterozygous or homozygous rare (minor allele frequency <0.01) variants in BM zone genes (**Fig. 5d** and **Supplementary table 15**). Odds ratio (OR) analysis revealed that homozygous pLoF variants in 41 BM zone genes were significantly enriched in affected individuals (OR=3.4, 95%CI=2.1-5.9, *p*<0.001; **Fig. 5e**, **Supplementary table 16**, and **Supplementary Fig. 7a**). Notably, homozygous pLoF variants in collagens and glycoproteins showed increased ORs (**Supplementary Fig. 7b**), mirroring the disease phenotype analysis (**Fig. 5c**). We next stratified the BM zone genes by known disease association evidence and found the OR for homozygous pLoF variants in high evidence genes was greatly elevated for affected 100KGP individuals (OR=9.8, 95%CI=4.0-33.0, *p*<0.001; Fig. **5e**). We also observed a trending OR increase for homozygous pLoF variants in genes with moderate to no disease associations (OR=1.5, 95%CI=0.8-3.1; **Fig. 5e**), suggesting that these could be new drivers of human disease associated with BM pathology.

### Disease-associated variants in *MATN2* perturb secretion and translation

Among the 41 homozygous pLoF variant genes (**Supplementary table 16**), 29 fitted a high penetrance variant filtering strategy (**Methods**). Notably, 11 of these genes had limited or no known disease assocations and we focused on three of these that appeared to be possible candidates underlying BM disease (**Fig. 5f**). First, we examined variants in *MATN2*, as three families were identified carrying unique biallelic *MATN2* variants with a range of multisystem phenotypes (**Fig. 6a**, **Supplementary notes**, and **Supplementary Fig. 8a**). The matrilin MATN2 is a poorly understood matrix protein that has not been linked to human genetic disease. To determine whether these variants disrupt MATN2 translation and secretion, we used human podocytes that secrete endogenous MATN2 and assemble a BM-like ECM^47^. CRISPR-Cas9 knockdown of *MATN2* nearly abolished the ECM fraction of MATN2, which was rescued by over-expression of wild-type V5-tagged *MATN2* (**Fig. 6b** and **Supplementary Fig. 8b**). In contrast, over-expression of the *MATN2^c.746G>C, p.Cys249Ser^-V5* missense variant resulted in MATN2 accumulation in the cellular fraction, suggesting a defect in secretion (**Fig. 6b**). Structural modelling showed disruption of a disulfide bond in the EGF1 domain of MATN2^c.746G>C, p.Cys249Ser^, consistent with a misfolded protein (**Supplementary Fig. 8c**). In a similar rescue experiment the MATN2^c.1585del, p.Cys529Valfs*13^-V5 frameshift variant protein was not detected in the cellular or ECM fractions, implying defective translation (**Fig. 6b** and **Supplementary Fig. 8d**). Additionally, a minigene splicing assay^48^ confirmed aberrant splicing and the introduction of a premature stop codon for the predicted splicing variant *MATN2*^c.1082+3-1081+6del^ (**Fig. 6c** and **Supplementary Fig. 8e**). These analyses suggest that the *MATN2* variants perturb proper protein production, thus validating our variant filtering strategies. As *C. elegans* and zebrafish lack *MATN2* orthologs, we next analyzed podocyte-derived matrix by proteomics to determine whether loss of *MATN2* affects BM composition. Importantly, levels of core BM components nidogen and collagen IV were decreased, laminin chains were variably affected, and perlecan was increased (**Fig. 6d** and **Supplementary table 17**). Together, these studies establish that new LoF *MATN2* variants could be associated with human disease and that loss of *MATN2* might alter BM composition.

**Figure 6.**
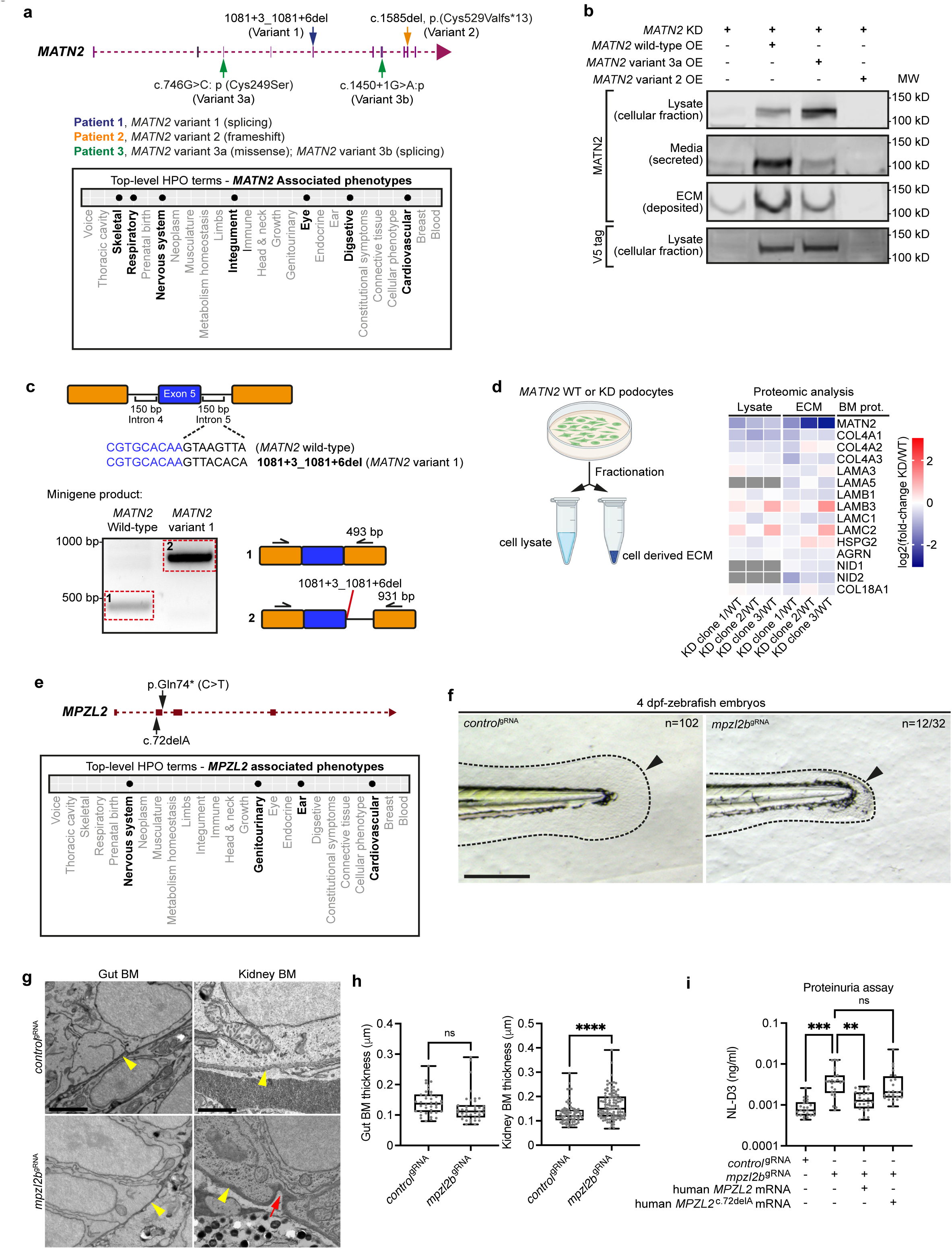
New loss-of-function *MATN2* variants are associated with human disease and MPZL2 loss disrupts zebrafish basement membranes. **a**, *MATN2* genomic structure indicating four pLoF variants (top) and associated phenotypic abnormalities represented as HPO terms (bottom, bolded). **b**, Western blots of MATN2 in lysate and ECM fractions derived from endogenous MATN2-depleted human podocytes over-expressing V5-tagged wild-type or variant MATN2. **c**, *In vitro* minigene splicing assay (**Methods**) demonstrating altered splicing of *MATN2* variant 1. **d**, Fold change in BM component levels upon *MATN2* knockdown as determined by fractional proteomic analysis of podocyte-derived ECM. **e**, *MPZL2* genomic structure indicating two 100KGP pLoF variants (top) and associated phenotypic abnormalities (bottom, bolded). **f**, Brightfield images of tail regions (dashed lines) in *control*^gRNA^ and *mpzl2b*^gRNA^-injected 4-dpf zebrafish embryos. Arrowheads highlight reduced fin fold extension in *mpzl2b* crispants. Scale bar represents 100 µm. **g**, Transmission electron microscopy (TEM) of gut and kidney BMs (yellow arrowheads) in control and *mpzl2b* crispant embryos. Red arrows indicate BM irregularities. Scale bar represents 2 µm (gut), 1 µm (kidney). **h**, Quantification of gut (n = 39 each) and kidney (n = 100 each) BM thickness . \*\*\*\**p*-value < 0.0001, ns—not significant; unpaired Student’s *t* test. **i**, Assessment of proteinuria (NL-D3 levels) in *mpzl2b* crispants injected with wild-type human *MPZL2* mRNA or *MPZL2*^c.72del^ variant mRNA (n = 24 each). \*\*\**p*-value < 0.001, \*\**p*-value < 0.01; ns—not significant; one-way ANOVA with post-hoc Dunnett’s test. For boxplots, edges indicate the 25^th^ and 75^th^ percentiles, the line in the box represents the median, and whiskers mark the minimum and maximum values.

### BM dysfunction linked to *MPZL2* and *LAMA5* variants

Next, we examined two genes with high penetrance variants but limited BM-associated disease phenotypes: *MPZL2* and *LAMA5* (**Fig. 5f**). MPZL2 is a transmembrane glycoprotein^49^ with genomic variants associated with hearing loss^50^. We identified one 100KGP individual carrying a homozygous pLoF variant *MPZL2*^c.72del^ with kidney cysts, cerebral aneurysms, and hypertension—phenotypes connected to BM defects (**Fig. 6e**, **Supplementary Fig. 8a**, and **Supplementary notes**). To investigate whether *MPZL2* loss affects BM structure or function, we CRISPR-depleted *mpzl2b* in zebrafish and observed abnormal fin development and thickened kidney BMs (**Fig. 6f-h**). Consistent with impaired BM function, we also observed elevated proteinuria (**Fig. 6i**). Notably, co-injection of human *MPZL2* mRNA rescued kidney function, while the *MPZL2^c.72del^* variant mRNA failed to rescue filtration defects (**Fig. 6i**). Together, these results indicate that MPZL2 regulates BM structure and function and suggest the *MPZL2*^c.72del^ variant could be a driver of BM-associated disease.

LAMA5 is a laminin alpha chain that together with LAMB2 and LAMC1 forms the laminin-521 network in BMs^51^. Its loss results in skin and kidney defects in mice^52^ and in humans biallelic missense variants are associated with phenotypes across multiple systems^53, 54^. We identified a family with two early fetal losses where both fetuses carried two pLoF *LAMA5* variants: a truncating variant *LAMA5*^c.9489C>T;p.Tyr3163*^ in trans to a predicted splice variant *LAMA5*^c.3282+5G>A^ (**Fig. 7a** and **Supplementary notes**). Minigene analysis confirmed altered splicing for *LAMA5*^c.3282+5G>A^ (**Fig. 7b**). Both fetuses had abnormalities in multiple systems, including absent or dysplastic kidneys (**Fig. 7a**, **Supplementary Fig. 8a**, and **Supplementary notes**). We acquired the 19-week fetal kidney tissue and observed multiple cysts and defective glomerular architecture (**Fig. 7c**). Visualization of the laminin-521 network with LAMB2 immunofluorescence revealed diffuse intracellular LAMB2 signal within abnormal glomeruli in the dysplastic kidney, which contrasted to crisp glomerular BM localization in a normal fetus (**Fig. 7d**). Furthermore, pan-laminin (LAM) immunofluorescence was dramatically thickened around abnormal structures in the dysplastic kidney, indicating that loss of *LAMA5* disrupts regulation of other BM laminin molecules (**Fig. 7d**). To further investigate the impact of *LAMA5* loss, we depleted *lama5* in zebrafish and observed abnormal fin fold development, ultrastructural irregularities and indentations in the gut and kidney BMs, and increased proteinuria (**Fig. 7e-g**). Together, our findings demonstrate *LAMA5* is crucial for vertebrate kidney BM structure and function and establish that new biallelic LoF variants in *LAMA5* cause an early onset human developmental disorder, thus providing a likely diagnosis for the family.

**Figure 7.**
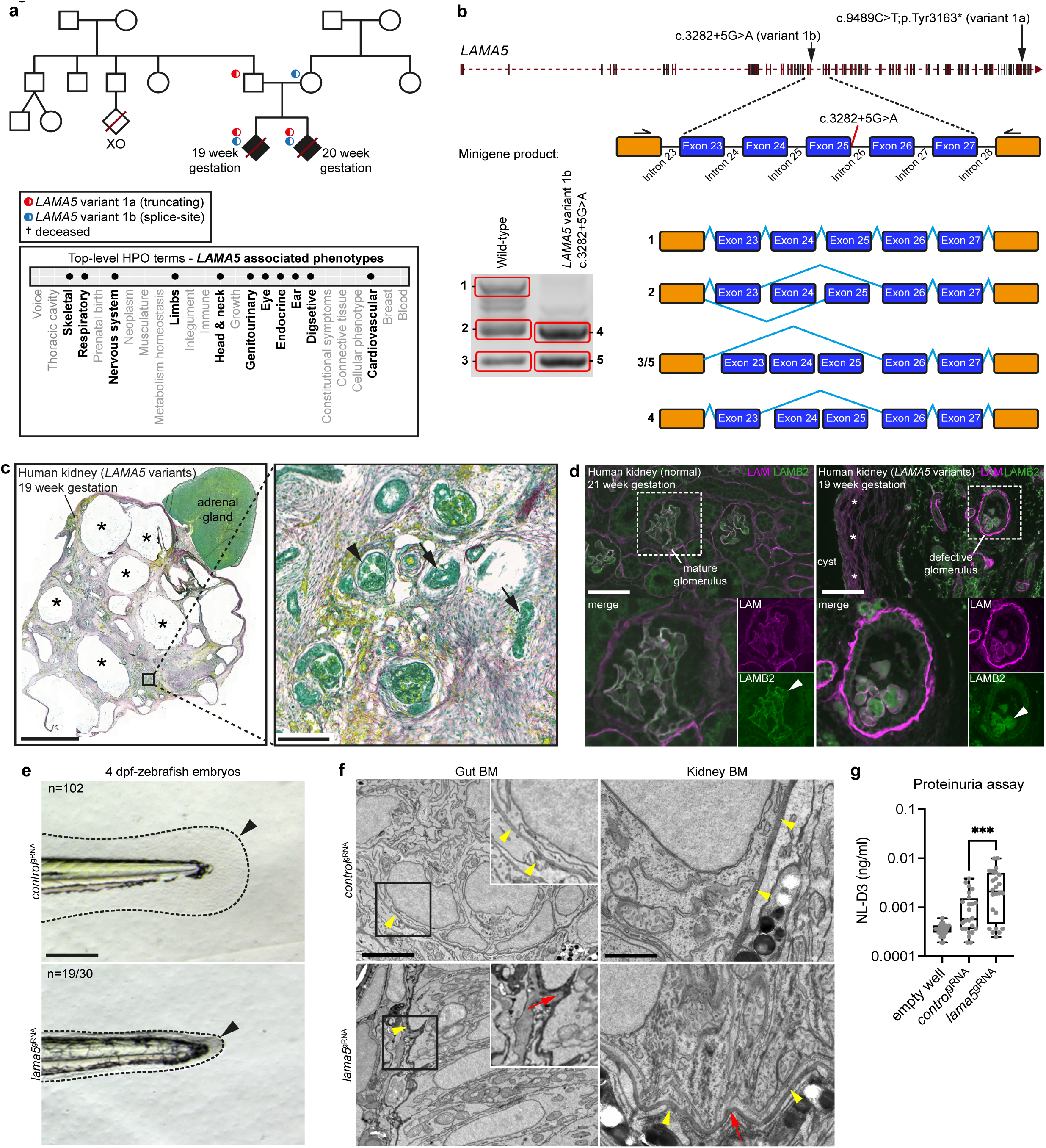
New genomic variants in *LAMA5* affect basement membrane structure and function. **a**, Top, genetic pedigree for a 100KGP family carrying two *LAMA5* pLoF variants. Bottom, phenotypic abnormalities observed in two *LAMA5* variant-carrying fetuses (19 and 20 week gestation) represented as bolded HPO terms. **b**, Top, *LAMA5* genomic structure and variant locations. Bottom, *in vitro* minigene splicing assay demonstrating altered splicing of *LAMA5* variant 1b. **c**, Left, Picrosirius red/fast green staining of the 19-week dysplastic kidney; asterisks indicate cysts. Boxed area is magnified on the right; arrowhead and arrows denote abnormal glomerulus and tubules, respectively. Scale bars represent 5 mm (left), 100 µm (right). **d**, Pan-laminin (LAM, magenta) and laminin β2 (LAMB2, green) immunofluorescence in wild-type (left) and dysplastic (right) fetal kidney sections. Asterisk indicates laminin surrounding a cyst. Boxed regions are magnified below. Arrowheads, see text. Scale bar represents 50 µm. **e**, Brightfield images of zebrafish tail regions (dashed lines) in *control*^gRNA^ and *lama5*^gRNA^-injected 4-dpf embryos. Arrowheads highlight reduced fin fold extension in *lama5* crispants. Scale bar represents 100 µm. **f**, TEM of gut and kidney BMs (yellow arrowheads) in control and *lama5* crispants. Boxed regions are magnified in insets. Red arrows indicate BM morphological irregularities. Scale bars represent 2 µm (gut), 1 µm (kidney). **g**, Proteinuria in control versus *lama5* crispants (n = 24 each). \*\*\**p*-value < 0.001, unpaired Student’s *t* test. For boxplots, edges indicate the 25^th^ and 75^th^ percentiles, the line in the box represents the median, and whiskers mark the minimum and maximum values.

## Discussion

We developed a pipeline combining bioinformatic approaches and *in vivo* localization to comprehensively annotate BM composition (**Fig. 8**). Our analyses define the makeup of the human BM zone: 160 BM matrix proteins and 62 cell surface interactors (CSIs). We show that 57 human BM proteins and 45 CSIs are shared with mice, zebrafish, *Drosophila*, and *C. elegans*. More than 50 are found solely in vertebrates and consist largely of proteoglycans, collagens, collagen modifying enzymes, and expanded ADAMTS protease family members. Using human transcriptomic and proteomic datasets, we identified tissue-level signatures of BM gene expression and composition, which could have clinical utility as organ health readouts and disease biomarkers. There were only five abundant and common BM zone proteins across analyzed tissues: laminin, collagen IV, collagen VI, nidogen, and perlecan. This suggests that a limited core BM scaffolding supports vast tissue-specific combinations of other BM residents, which likely provide tissues unique mechanical and signaling support and help account for the many functions of BMs.

**Figure 8.**
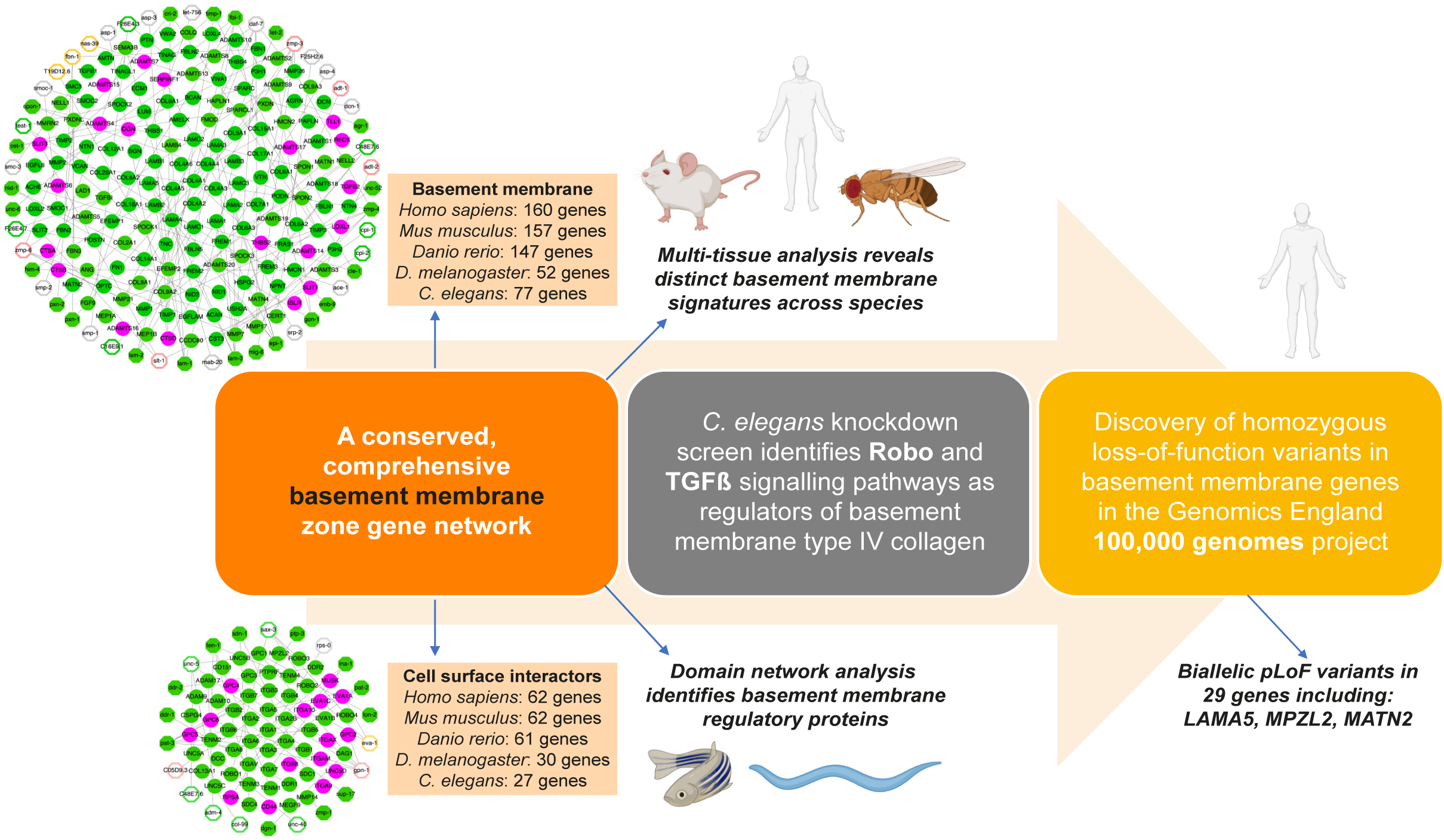
A basement membrane discovery pipeline. Summary of multidisciplinary approaches and model systems used in this study that established a comprehensive network of BM matrix proteins and their cell surface interactors, new regulators of BM structure and function, and identified BM component genomic variants associated with phenotypic abnormalities in 100KGP.

Biochemical experiments and studies of embryonic BM assembly have provided insights into how the core BM scaffold is first assembled^5, 10, 55–60^. BM regulation during tissue growth, shaping, remodeling, and homeostasis, however, is less well understood^61^. By constructing domain-based networks, we identified conserved candidate regulatory hub proteins, of which the heparan sulfate proteoglycan perlecan (HSPG2) shares the most domains with other BM zone proteins. We discovered a role for perlecan late in *C. elegans* larval development in maintaining gonadal tissue shape. Perlecan loss in *Drosophila* also results in tissue deformation^37, 38^, suggesting tissue sculpting is a general perlecan function. In addition, we discovered perlecan loss resulted in fibrillar and globular BM aggregates. Given perlecan’s many shared domains, it might help maintain tissue form in part by connecting BM constituents. Other BM regulatory hub candidates include signaling molecules, proteases, and heparan sulfate proteoglycans. Some of these have known post-embryonic roles in BM growth (*e.g.* GON-1 (ADAMTS9) and MIG-6 (PAPLN))^28, 62^ and we establish here a role for SAX-3/ROBO in controlling BM composition. Thus, network connectivity analysis provides a new approach to uncover key BM regulatory proteins.

To further investigate mechanisms of post-embryonic BM regulation, we performed a comprehensive RNAi screen of 77 BM zone genes on the growing *C. elegans* gonadal BM. Loss of ADT-2 (ADAMTS3) protease caused a sharp reduction in laminin and collagen IV levels and misshapened gonads. As *ADAMTS3* variants are associated with abnormally expanded lymphatic vessels in Hennekam syndrome^63^, ADAMTS3 might control tissue shape by regulating BM composition. We discovered many genes that function to restrict BM collagen IV levels. These collagen IV limiting mechanisms might break down during senescence and disease, as altered collagen IV levels is a hallmark of aging, diabetes, and Alzheimer’s disease^19, 64–66^. We found the TGFβ pathway and ROBO restrict collagen IV accumulation in *C. elegans* and zebrafish. Interestingly, collagen IV binds and regulates the ligands of these receptors^12, 67^. Thus, our findings indicate that interactions between these pathways and collagen IV are reciprocal and may be a mechanism to set and maintain collagen IV levels and buffer levels of signaling.

Using our BM zone gene network, we revealed human phenotype associations for more than 100 genes that affect all major organ systems, thus consolidating the growing link between BM genes and human disease^14^. Through genomic analyses of the 100KGP rare disease cohort, we also identified high penetrance homozygous pLoF variants in 11 BM zone genes with limited or no previous associations to BM pathology and human disease, and we characterized *MATN2*, *MPZL2,* and *LAMA5* variants. We report a new disease association for MATN2 and demonstrate that LoF variants in *MATN2* impact translation, secretion, and splicing. Biochemical analyses suggest MATN2 interacts with many BM proteins^68^, and we found that depletion of MATN2 affected levels of core BM components in human podocyte-derived ECM. MPZL2 and LAMA5 are BM zone proteins with limited connections to disease and BM pathology. A LoF variant in the adhesion protein MPZL2 (*MPZL2^c^*^.72delA^) was previously associated with hearing loss^50^, and we identified a 100KGP patient with additional nervous system, cardiovascular, and kidney defects, thus extending the phenotypic spectrum of *MPZL2^c^*^.72delA^. We used the zebrafish kidney to assay BM structure and BM filtration function and found that disruption of MPZL2 resulted in thicker, irregular kidney BMs and kidney filtration defects. MPZL2 is localized to the ear BM zone^50^, and it is likely that similar BM defects could lead to sensorineural hearing loss. Consistent with this notion, collagen IV (*COL4A3/4/5*) variants disrupt BMs and are also associated with hearing impairment^69^. *LAMA5* is a laminin alpha chain whose knockout is embryonic lethal in mice^70^. Two missense variants *LAMA5*^V3140M^ and *LAMA5*^R286L^ are associated with a range of human phenotypes but have not been connected to BM pathology^53, 54^. We identified new LoF *LAMA5* variants linked to human fetal lethality and defects in kidney BMs and kidney formation. Depletion of *lama5* in zebrafish also resulted in irregular BM morphology and BM function in the kidney. Together these findings provide compelling evidence that variants in *MATN2*, *MPZL2,* and *LAMA5* could lead to BM pathology in humans.

The network of more than 200 BM-associated components defined in this study provide BMs astonishing complexity, which likely underlies the diversity of BM functions that are vital to human health. The multi-disciplinary investigative pipeline established here to identify BM components, sites of localization, key BM regulatory nodes, and disease association provides a framework for discovering mechanisms that dictate BM function and regulation. We expect this will ultimately facilitate earlier disease detection, improve prognosis prediction, and inform new therapies for BM-associated human disease.

## Methods

### Identification of basement membrane and cell surface interactor genes

To build a comprehensive BM zone network of genes encoding BM and CSI components, we adopted the following strategy: (i) initial identification of BM zone candidates via gene ontology, (ii) expansion of the BM gene network, and (iii) verification of BM zone localization for candidates (detailed in **Supplementary notes**). Briefly, to obtain a primary list of candidate genes, we used Gene Ontology Resource (release 2020-07^30, 31^) to search for human genes catalogued under the ‘Basement membrane’ cellular component term (GO:0008003/GO:0005605) and retrieved 103 genes. Next, we applied a network expansion strategy to include new candidate genes through 1) data curation, 2) analysis of BM expression signature in *C. elegans*, 3) conservation and enrichment analysis for BM protein domains, 4) identification of nearest BM interacting neighbors, and 5) endogenous fluorescent tagging of BM zone candidates in *C. elegans* (**Supplementary tables 1-7**). This expansion strategy added 160 candidates to our network. Next, we confirmed protein localization for 188 out of 263 total candidates based on published immunolocalization studies in human, rodents, zebrafish, *Drosophila*, and *C. elegans* (**Supplementary table 7**), and fluorescent protein tagging studies in *C. elegans* in this work (**Supplementary table 5**). We predicted localization to the BM zone for 34 candidates based on evidence for protein-protein interaction with other BM/CSI candidates and BM substrate-cleaving activity for proteases (**Supplementary Fig. 1a**). We further categorized 40 candidates for which we could not confirm or predict BM zone localization under “insufficient evidence” and identified 16 GO mis-annotated genes, all of which were removed, and the final integrated verified BM zone network comprised 160 BM components and 62 CSIs. We adopted a colour code to differentiate the localization evidence for the components in our network: green for confirmed localization to BM zone; magenta for predicted to localize to the BM zone; and grey for insufficient localization evidence. To curate this list, we created an open access, interactive resource: ***b***asement ***m***embrane**Base** (https://bmbase.manchester.ac.uk/), which details background, BM gene lists, localization data, component descriptions, human disease descriptions, experimental systems (*C. elegans*, *Drosophila*, zebrafish, mouse, human), reagents, and protocols.

### Hierarchical clustering of gene expression and variance analysis

Baseline gene expression datasets screened for multi-tissue and non-disease gene expression, were downloaded from Expression Atlas^71^ (www.ebi.ac.uk/gxa/, release 31, for both human and mouse (see **Supplementary table 9**). Tissue expression profiles of the BM and CSI genes were clustered using hierarchical clustering on Spearman distance in R (v3.6.4). We calculated gene expression variance across all tissue assays using R, and binned BM and CSI genes into groups based on variance quartiles (low, medium, high, and very high; see **Supplementary table 10**).

### Basement membrane domain-based network analysis

To construct a domain-based BM network, we mapped the complement of putative BM protein domains obtained in the domain enrichment analysis (see **Supplementary notes** and **Supplementary table 3**) onto our verified BM zone gene network using Cytoscape^72^ v.3.8.1 where nodes in the network represent BM zone proteins and a single line connecting two nodes represent a set of BM domains common to both. We used the NetworkAnalyzer plugin for Cytoscape v.2.8^33^ to compute two network topological parameters to reveal potential organization and interaction of BM zone proteins—*network degree* and *betweenness centrality*. The network degree score measures the number of connections for a given node (i.e., lines radiating out of the node) and indicates the number of proteins in the network that share one or more domains with a given protein. Biologically, a high network degree score could indicate the different roles of a BM zone protein. The betweenness centrality score measures how central a given node is, based on the extent of shortest paths across the network that pass through that node. Thus, a protein with a high betweenness centrality score may contain domains typically found in distinct sub-networks, suggesting that the protein could bridge interactions between diverse groups of BM zone proteins.

### *C. elegans* and zebrafish strains

*C. elegans* and zebrafish strains used in this study are listed in **Supplementary table 18**. Worms were reared at 20°C on nematode growth medium plates seeded with OP50 *Escherichia coli* according to standard procedures^73^. Zebrafish were maintained and staged according to established protocols^74^ and in accordance with the project license P1AE9A736 under the current guidelines of the UK Animals Act 1986. Embryos were collected from group-wise matings of y-crystallin:mcherry/l-fabp10:NL-D3 fish.

### CRISPR-Cas9 endogenous fluorescent tagging in *C. elegans*

To generate endogenous mNeonGreen or mRuby2 tags for 17 BM and 8 CSI candidate genes in *C. elegans*, we used CRISPR-Cas9-mediated genome editing with a self-excising hygromycin selection cassette as described previously^4, 28^. Position of tags and fluorophores used for each genomic locus are detailed in **Supplementary Fig. 2**.

### RNAi in C. elegans

All RNAi constructs were obtained from the Vidal^75^ and Ahringer^76^ libraries, except for the following clones: *sax-3*, *zmp-4*, *cpi-2*, *Y64G10A.7*, *daf-1*, *tig-3*, *daf-7*, *sma-6*, *smp-2*, and *mab-20*, which were constructed from PCR fragments corresponding to the longest transcripts of the respective genes, as described previously^4^. RNAi experiments were performed using the feeding method^77^ and according to protocols detailed in Jayadev et al., 2019^4^. RNAi was initiated in synchronized L1 larvae, and animals were fed for 24-72 h at 20°C depending on the experiment. For the RNAi screen, we first assessed laminin (LAM- 2::mNG) and type IV collagen (EMB-9::mRuby2) gonadal BM fluorescence levels visually. Quantifications were then performed for all knockdown conditions other than positive controls where visible changes in fluorescence intensity was observed. At least 20 animals each were examined for both visual and quantitative assessments of BM fluorescence.

### CRISPR-Cas9 knockdown in zebrafish

CRISPR-Cas9 knockdown was performed as previously described^40^. Four gRNAs for each gene (**Supplementary table 19**) were used to generate crispant embryos with knockout-like phenotypes in the F_0_ generation. gRNAs (Merck) were resuspended to 20 µM and aliquoted for storage. On the day of injection, the RNP injection mix [5 µM EnGen^®^ Spy Cas9 NLS (NEB Cat# M0646T) mixed with 2.5 µM of each gRNA and Cas9 buffer] was freshly prepared, and 5.0 nl was injected into one-cell stage wild type zebrafish embryos, which were then maintained and processed for further analyses when they reached the required stage of development.

### Kidney filtration assay in zebrafish

The *NL:D3* transgenic zebrafish^78^ were injected at the one-cell stage with gRNAs targeting *adamts3* or *robo1* (**Supplementary table 19**) and Cas9 protein. Crispant *NL:D3* embryos were grown to 4 days post-fertilization (dpf) and then placed in wells of a 96-well plate. Three embryos were placed in a single well and 12 wells were set up per experimental sample. Embryo media (E3) was removed and replaced with 200 µl fresh E3. These embryos were then left for 24 h before 50 µl of E3 was extracted from each well and placed into corresponding wells of an opaque 96-well plate. 50 µl of Nano-Glo® Luciferase reporter buffer and substrate (Promega #N1120) was added to the cultured E3 media. The 96-well plate was spun at 700 rpm for 1 min before luminescence intensity was determined on a Flexstation 3 multi-mode microplate reader (Molecular Devices).

### TGFBR1 inhibition

Zebrafish embryos were grown to 24 hours post-fertilization (hpf) and manually dechorionated. Embryos were then placed in 50 µM SB431542 (Calbiochem, #616461) or 0.1% DMSO vehicle control and cultured for a further 48 h with a change in media (containing drug or vehicle) after 24 h. Embryos were fixed in 4% PFA at 72 hpf.

### Human tissue samples

Postmortem fetal tissue from the *LAMA5* cases was used with prior parental consent for use in research. The control 21-week human fetal kidney sections were acquired in a previous study^79^ and were initially provided by the Joint MRC/Wellcome Trust Human Developmental Biology Resource (HDBR) (http://hdbr.org). We were unable to obtain further clinical details or tissue samples from the reported individuals with *MPZL2* and *MATN2* variants.

### Histological staining and immunofluorescence

Formalin-fixed paraffin-embedded (FFPE) human fetal kidney sections were dewaxed using xylene and hydrated using a graded series of ethanol solutions (100%-70%) to water. The slides were then stained using Picrosirius red with Fast green counterstain for 1 hour and washed with 1% acetic acid. Slides were then dehydrated using ethanol and xylene before coverslipping.

For pan-laminin and LAMB2 immunofluorescence, FFPE human fetal kidney sections were dewaxed, hydrated, and heated in 10 mM sodium citrate buffer (pH 9.0) for 15 min for antigen retrieval. After blocking with 1% BSA (prepared in PBS) for 1 h at room temperature, sections were stained with a rabbit anti-laminin (Abcam, ab11575; 1:200) and mouse anti- LAMB2 (Millipore, MAB2066; 1:50) antibodies overnight at 4°C. For MATN2 immunofluorescence, human podocytes were fixed in 4% PFA and stained using a rabbit anti-MATN2 (ProteinTech Ltd, 24064-1-AP; 1:150) and a mouse anti-V5 (Abcam, ab2767; 1:150). For type IV collagen immunofluorescence, 5-dpf zebrafish embryos were fixed in 4% PFA overnight at 4°C and embedded in OCT. 5-μm-thick cryosections were air-dried, permeabilized with acetone at -20°C for 8 minutes, blocked with 10% FBS and 5% donkey serum in PBS-Triton X for 1 h at room temperature, and stained with a rabbit anti-type IV collagen (Abcam, ab6586; 1:250) overnight at 4°C.

Species-specific secondary antibodies conjugated with either Alexa-Fluor 488 (Invitrogen Antibodies, A11008; 1:400) Alexa-Fluor 594 (Invitrogen Antibodies, A21203; 1:400) or Alexa-Fluor 647 (Invitrogen Antibodies, A-21235) were used, and the slides mounted with Prolong Diamond antifade (ThermoFisher, P36961).

### Microscopy and Image Processing

For *C. elegans* experiments, fluorescence images were acquired at 20°C on an AxioImager A1 microscope (Carl Zeiss) controlled by μManager software^80^ and equipped with an electronmultiplying charge-coupled device camera (Hamamatsu Photonics), a 40× Plan-Apochromat (1.4-NA) objective, a spinning disc confocal scan head (CSU-10; Yokogawa Electric Corp.), and 488- and 561-nm laser lines. Worms were mounted on 5% noble agar pads containing 0.01 M sodium azide for imaging. For the BM zone gene tagging experiments and the RNAi screen, we captured single-slice images at the middle focal plane of animals where most or all the gonadal and pharyngeal tissue cross sections were in focus. For the perlecan knockdown experiments, we acquired z-stacks at 0.37-μm intervals spanning the surface of the gonad and generated sum projections. All quantifications of mean fluorescence intensity were done on raw images in Fiji v.2.0^81^. We drew ȣ30 pixel-long line scans along the BM to obtain raw values of mean fluorescence intensity. Background intensity values were obtained by averaging two-lines scans of similar length in regions adjacent to the BM with no visible fluorescence signal.

For zebrafish experiments, images were collected on a Leica TCS SP8 AOBS inverted confocal using a 20x / 0.50 Plan Fluotar objective and 3x confocal zoom. The confocal settings were as follows, pinhole 1 airy unit, scan speed 1000Hz unidirectional, format 1024 x 1024. Images were collected using hybrid detectors with the following detection mirror settings; AlexaFluor 494-530nm using the white light laser with 488nm (10%) laser line respectively. For quantification of fluorescence, we obtained raw integrated density measurements for the whole field of view in Fiji. Brightfield images of zebrafish embryos were acquired using a Leica M205 FA upright stereofluorescence microscope. For electron microscopy, samples were prepared according to protocols described previously^82^. Images were taken on T12 Biotwin transmission electron microscope. Distances were measured in Fiji/ImageJ using grid method measurements were taken and normalized to the length of the glomerular basement membrane.

For human histological staining and immunofluorescence, slides were imaged using a 3D- Histech Pannoramic-250 microscope slide-scanner. Fluorescence images were acquired using a PCO.edge camera with a 20x/0.80 Plan Apochromat objective (Zeiss) and the FITC and Texas Red filter sets. Transmitted light images were acquired using a CIS VCC FC60FR19CL camera with a 20x/0.80 Plan Apochromat objective (Zeiss). Snapshots of the slide-scans were taken using the Case Viewer software version 2.4.0.119028 (3D-Histech). slides were imaged using a 3D Histec Pannoramic250 slide scanner microscopy. For human podocyte experiments, fluorescence images were collected on a *Zeiss Axioimager.D2* upright microscope using a *40x / 0.75 EC Plan-neofluar* objective and captured using a Coolsnap HQ2 camera (Photometrics) through Micromanager software v1.4.23. Specific band pass filter sets for *DAPI, FITC and Far red (Alexa 647)* were used to prevent bleed through from one channel to the next. Images were then processed and analyzed using *Fiji ImageJ* (version 1.5.2; National Institutes of Health, Bethesda, MD, USA).

### Human disease phenotypes

We manually searched for each of the BM and CSI genes within the following databases: Human Phenotype Ontology (HPO) database^44^ (https://hpo.jax.org/app/), the Genomics England PanelApp^45^ (https://panelapp.genomicsengland.co.uk), and the Online Mendelian Inheritance in Man (OMIM) database (McKusick-Nathans Institute of Genetic Medicine, Johns Hopkins University Baltimore, MD; https://omim.org/). Top-level HPO terms associated with each OMIM disease entry listed were grouped under individual BM genes.

### Genomic analysis

We screened whole genome sequencing data from the Genomics England 100,000 genomes project (100KGP)^29^ for variants in the verified BM zone gene network. The project was formally registered and approved (Genomics England Research Registry, RR320). We extracted all BM and CSI genomic variants from an aggregated set of variants available for 63,039 individuals recruited to the rare disease arm of the 100KGP; this data is comprised of 34,842 individuals affected with a rare disease across broad disease areas and 28,197 unaffected relatives at the time of recruitment. The minor allele frequency was calculated from Genomics England aggregated variant calling format and pLoF variants were identified after annotation through Ensembl variant effect predictor to a predefined list of transcripts for the BM and CSI gene network. Variants matching one or more of the following criteria were included: stop_lost, start_lost, stop_gained, transcript_ablation, splice_acceptor_variant, splice_donor_variant, frameshift_variant. Gene burden testing was performed using Test Rare vAriants with Public Data (TRAPD)^83^ and identified only modest signals for previously known disease-causing genes under dominant and recessive models (data not shown). Due to the heterogeneous nature of the cohort, we utilized a logical filtering strategy to identify rare/novel pLoF variants enriched in affected individuals under the assumption of high penetrance: ≤1 homozygous pLoF variant impacting canonical transcript in gnomAD v2.1 with the high confidence LoF flag, and no individuals with homozygous pLoF variants in unaffected 100KGP cohort; genes with multiple affected individuals were prioritized. Genes fitting these criteria were further evaluated for individuals carrying heterozygous pLoF variants *in-trans* to another potential damaging mutation; these were highlighted through Exomiser^84^ results available in the Genomics England research embassy. Statistical analyses and graphics were created in R software. We report odds ratios and *p-*values from the fisher exact test using the oddsratio function in the ‘epitools’ package for R. Case histories for individuals carrying *MATN2*, *MPZL2*, or *LAMA5* variants are provided in **Supplementary notes**.

### Minigene splicing assay

Minigene vectors for both wild-type and variant sequences for *MATN2* and *LAMA5* were assembled using the SK3 minigene vector (a derivative of the pSpliceExpress minigene splice reporter vector, gifted from Stefan Stamm, Addgene #32485) as previously described^48^. HEK293 cells were cultured to 40-60% confluency in 2ml of Dulbecco’s modified Eagle’s medium high-glucose, DMEM (Sigma), supplemented with 10% foetal bovine serum (Sigma) in a 6-well cell-culture plate at 37°C with 5% CO_2_. Cells were transiently transfected with 2.5μg of minigene vector using Lipofectamine 3000/LXT (Thermofisher Scientific) using manufacturer’s protocol. Following 18h incubation at 37°C with 5% CO_2_, RNA was extracted using 1 ml of TRI-Reagent® (Sigma) per well and further purified using an RNAeasy cleanup kit (Qiagen). An equal amount of RNA for each sample was converted to cDNA using Primescript^TM^ RT Reagent Kit (Takara Bio), or Superscript IV (Thermofisher Scientific). cDNA was amplified using Q5 (NEB) or KOD polymerase and specific minigene primers (**Supplementary table 20**). Finally, PCR products were run on an agarose gel (1-4%) supplemented with SybrSafe (Thermofisher Scientific) for visualisation on a UV transilluminator. PCR products were purified using QIAquick PCR purification kit (Qiagen) and sequenced by Sanger sequencing (Eurofins Genomics).

### 3D structure prediction

The MATN2 (O00339) protein sequence was obtained from the UniprotKB database. EGF1 domain (aa: 238-278) sequences of either the wild-type MATN2 or MATN2 variant 3a were submitted to the Phyre2 web portal to generate three dimensional models^85^. Templates (c1lpkA for MATN2_EGF1) were selected based on maximum identity and query coverage. Visualization of the variant EGF domain was performed using PyMOL (The PyMOL Molecular Graphics System, Version 1.2r3pre, Schrödinger, LLC).

### *MATN2* gene knock-down and rescue

A *MATN2* knock-down line was generated in human immortalised podocytes^86^ by CRISPR- Cas9. Cas9 expressing podocytes were established by transduction with pLentiCas9_v2 (addgene plasmid 52961) followed by Puromycin selection. These Cas9-podocytes were transfected twice with two sgRNA targeting *MATN2* (5’-GTCACGATCATTATGACCCG-3’; 5’-CTTGACCTTTGCATAGTCAT-3’; Merck) using RNAiMAX (Thermofisher Scientific) in a 6- well plate, and depletion of MATN2 confirmed by western blot. Lentiviral vectors expressing MATN2 coding sequences (wild-type, Variant 2 or 3a) under the control of EF1a promoter were designed, cloned, and packaged (Vectorbuilder, vectors will be made available through Addgene). 10^6 MATN2-KD podocytes were seeded in 10 cm dishes and transduced; briefly, to 964 µl of serum free media, 16 µl polybrene (5 mg/ml stock) and 20 µl of 3.78 x10^8 TU/ml virus were added and incubated for 6 hours. The virus was then removed, cells washed, and fresh complete RPMI media added. Stably transduced cells were differentiated for 8 days before harvesting cell lysate, media, and deposited ECM. Western blotting was performed using 4–12% BisTris gradient gels with the MES buffer system according to the manufacturer’s instructions (Life Sciences). Reduced protein samples were transferred to nitrocellulose membranes then blotted using anti-MATN2 at 1:1000 dilution (ProteinTech Ltd, 24064-1-AP) with anti-rabbit 680 conjugate (Jackson Labs; 1:10000) as a secondary antibody. Proteins bands were visualized using an Odyssey Imaging System (LI-COR) at 700 nm. For PCR amplification of the V5 tag cDNA was extracted from cells and a PCR was performed using the QIAquick PCR purification kit (QIAGEN) and the following primer sequences: Forward- 5’ GCCTATCCCTAACCCTCTCC 3’ and Reverse- 5’ CATTCTTGACAGTGCTGCCA 3’.

### Sample preparation for mass spectrometry

Three clones were generated from a pool of a human podocyte cells that were knocked down for *MATN2*. The cells were then diluted and plated up in 96-wells plate to obtain colonies from a single cell (=pure population of cells with the same phenotype, i.e., MATN2- KD). Human immortalized podocytes^86^ (wild-type and the 3 MATN2-KD clones) were cultured for 14 days at 37°C in monoculture on uncoated 10-cm dishes using RPMI-1640 medium (Sigma-Aldrich, R-8758) supplemented with 10% fetal calf serum (Life Technologies), 1% insulin, transferrin, and selenium (Sigma-Aldrich, I-1184) and 1% penicillin-streptomycin. Samples were enriched for ECM proteins as previously described^47^. Briefly, cells and matrix were scraped into lysis buffer (10 mM Tris, 150 mM NaCl, 1% Triton X-100, 25mM EDTA, protease inhibitor) and incubated for 1 h followed by centrifugation at 14,000×g for 10 min. The supernatant (lysate) was kept and added with 5% SDS with 50 mM triethylamonium bicarbonate (TEAB). The pellet was resuspended in alkaline detergent buffer (20 mM NH4OH and 0.5% Triton X-100 in PBS) for 30 min to disrupt cell-matrix bonds. Following centrifugation, the remaining pellet enriched for ECM proteins was resuspended in 5% SDS with 50 mM TEAB.

The lysate and the ECM-rich fractions were processed for in-solution digestion as described previously^79^. To extract proteins, samples were lysed using a Covaris LE220+ Focused Ultrasonicator (Covaris). The protein samples were reduced with 5mM dithiothreitol for 10 minutes at 60°C and alkylated with 15mM iodoacetamide for 30 min in the dark. Protein concentration was quantified using a Direct Detect Spectrometer (Millipore, DDHW00010- WW), and 40 μg of proteins/sample was loaded onto S-trap columns (Protifi) and digested overnight with 1 μg sequencing grade modified trypsin (Promega, V5111) at 37°C. Peptides were eluted from the column with 30% acetonitrile and 0.1% formic acid aqueous solution, and collected by centrifugation at 4,000×g for 2 min. After desalting, the peptide samples were dried to completeness by vacuum centrifugation and submitted to analysis by mass spectrometry.

### Mass spectrometry data acquisition and analysis

Samples were analyzed by liquid chromatography-tandem mass spectrometry using a Thermo Exploris 480 mass spectrometer (Thermo Fisher Scientific). MS analysis was carried out using Proteome Discoverer v.2.4 SP1 (Thermo Fisher Scientific). Raw spectra data were searched against human SwissProt and TrEBML (v.2017-10-25) using the SEQUEST HT search tool. Tryptic peptides with up to 2 missed cleavage sites, and mass tolerance of 10 ppm for precursor ions and 0.02 Da for fragment ions were set for the search. Carbamidomethylation of cysteine was selected as fixed modification, and oxidation of proline, lysine and methionine, and N-terminal acetylation as dynamic modifications. Protein false discovery rate was set to 1%, and protein validation was performed using Target/Decoy strategy. Protein abundances were determined by label-free quantification based on precursor ion intensity. Results were filtered for significant FDR (≥0.1) master proteins identified with ≥ 1 unique peptide. The MS proteomics data have been deposited to the ProteomeXchange Consortium via the PRIDE partner repository^87^ with the dataset identifier PXD029266.

### Statistical analysis

For *C. elegans* and zebrafish experiments, statistical analysis was performed in GraphPad Prism v.7. Sample sizes were validated a posteriori through assessments of normality by log-transforming all datasets and then using the Shapiro–Wilk test. For comparisons of mean fluorescence intensities between two populations, we used an unpaired two-tailed Student’s t test (with Welch’s correction in cases of unequal variance between samples). To compare mean fluorescence intensities between three or more populations, we performed one-way ANOVA followed by a post hoc Dunnett’s multiple comparison test. For odds ratio comparisons in genomic analyses, we performed the Fisher’s Exact Test in R software. Boxplots were prepared in GraphPad Prism 7. Figure legends indicate sample sizes (n), statistical tests used, and *p-*values.

## Acknowledgements

This work was supported by a Wellcome Senior Fellowship award (202860/Z/16/Z) to R.L. and the National Institutes of Health grants R35GM118049 and R21OD028766 to D.R.S. M.R.P.T.M. was supported by the Global Challenge Research Fellowship program, E.H. by 129351-PF-16-024-01-CSM from the American Cancer Society; M.F. by a Kidney Research UK grant (RP_040_20180306), R.T.O. and H.T. by a Biotechnology and Biological Sceinces Research Council award (BB/S00047X/1), and D.S. by a National Institute of Child Health and Human Development award (1R01HD103805-01). The authors also acknowledge core funding from the Wellcome Trust (203128/Z/16/Z) to the Wellcome Centre for Cell-Matrix Research at the University of Manchester, and funding for equipment from the Biotechnology and Biological Sciences Research Council and the University of Manchester Strategic Fund. We thank Sara Payne, William Ramos-Lewis, and Liam Lewis for assistance with molecular biology; Peter Breen and Gary Ruvkun for sharing DAF-4::GFP translational reporter strains; staff from the Biomolecular Analysis, Bioimaging, and Electron Microscopy Core Facilities for advice and assistance; and Laura Kelley for helpful discussions. This research was made possible through access to the data and findings generated by the 100,000 Genomes Project. The 100,000 Genomes Project is managed by Genomics England Limited (a wholly owned company of the Department of Health and Social Care). The 100,000 Genomes Project is funded by the National Institute for Health Research and NHS England. The Wellcome Trust, Cancer Research UK and the Medical Research Council have also funded research infrastructure. The 100,000 Genomes Project uses data provided by patients and collected by the National Health Service as part of their care and support.

## Author contributions

Study conceptualization and experimental design: RJ, MRPTM, JME, DRS, RL; Experimental procedures, data acquisition and analysis: RJ, MRPTM, JME, SS, RWN, CL, AL, JFI, EH, QC, MF, NMK, HBT, EW, DRS, RL; Manuscript preparation: RJ, MRPTM, JME, DRS, RL; Manuscript editing and review: RJ, MRPTM, JME, SS, RWN, CL, AL, JFI, EH, QC, MF, NMK, HBT, RTO, EW, AA, HMS, SB, DS, DRS, RL; Genomic data availability: Genomic England Research Consortium; Genomic data analysis: JME, DS; Provided access to patient samples: HMS.

## Competing interests

The authors declare no conflicts of interests.

## Data availability statement

Primary data from the 100,000 Genomes Project, which are held in a secure Research Environment, are available to registered users (see https://www.genomicsengland.co.uk/about-gecip/for-gecip-members/data-and-data-access for further information). Immunoblotting source data are hosted at https://doi.org/10.6084/m9.figshare.c.5662348. All other data are provided in Supplementary tables.

## Supplementary Figure Legends

**Supplementary Figure 1. Verification strategies for basement membrane zone localization and conservation of network genes. a**, Venn diagrams illustrate how the localization of human proteins in the integrated basement membrane (BM) zone network was verified. The asterisk indicates a group of proteins with predicted localization whose *C. elegans* orthologs were detected in the BM zone through fluorescent tagging. **b**, Human BM matrix and cell surface interactor (CSI) genes are shown in boxes and the presence of corresponding orthologs in mouse, zebrafish, *D. melanogaster*, and *C. elegans* (**Supplementary Fig. 3** and **Supplementary table 7**) is represented in Venn diagrams. Animal illustrations made with <u>Biorender</u>.

**Supplementary Figure 2.** Visualization of basement membrane zone candidate localization in *C. elegans* through fluorescent protein insertions at endogenous gene loci. a, Worm gene schematics (adapted from https://wormbase.org) indicate genomic locations where mNeonGreen (green box), two tandem mNeonGreen (two green boxes), or mRuby2 (red box) fluorophores were inserted for BM and CSI candidates tagged in this study. For each gene, sgRNA, PAM site, and insertion site sequences are provided. Every gene was tagged at a single locus, except for *col-99*, which was tagged at two separate loci. Note that exons are colored magenta or cyan depending on alignment to the forward or reverse strand of the reference genome, respectively. b, Confocal middle-plane z-slices of tagged candidates in adult animals showing BM zone localization (yellow arrowheads) to either the pharyngeal or gonadal BM, except for FRAS/FREM/*C48E7.6*, which localizes to the body wall muscle BM. c, Confocal z-slices of candidates not detected in the BM zone but present in other matrices (blue arrowheads). Scale bar represents 25 μm.

**Supplementary Figure 3. Basement membrane zone gene families in *C. elegans*.** Diagrams illustrate the orthologous relationships of *C. elegans* genes and gene families (on the left) with human BM zone genes (on the right). Note the various one-to-many and many-to-many relationships between several genes for both species.

**Supplementary Figure 4. Expression variance of mammalian basement membrane zone genes**. Heatmaps indicate scaled gene expression of BM matrix and CSI genes across different (**a**) human and (**b**) mouse tissues (transcriptomic datasets used in this study detailed in **Supplementary table 9**). Genes are binned according to expression variance (low, moderate, high, very high; **Supplementary table 10**).

**Supplementary Figure 5.** Perlecan/UNC-52 and papilin/MIG-6 are hub proteins and depletion of *C. elegans* UNC-52 disrupts basement membrane organization. a, Conservation and enrichment of Interpro protein domains (as described in **Fig. 3a**) in verified CSI components. b, A domain-based interactome (as described in **Fig. 3b**) for *C. elegans* BM zone genes. c, Sub-network highlighting diverse groups of human BM zone proteins that share domains with papilin. d, Left, confocal sum projections of gonadal BM LAMC/LAM- 2::mNG in control and *unc-52* RNAi-treated early L3, young adult, or gravid adult animals (n = 10 animals examined each). Right, confocal middle-plane z-slices of HSPG2/UNC- 52::mNG, which is not detected at the gonadal BM (outlined in magenta) in the early L3 stage, but is visible in the young adult and gravid adult stages (yellow arrowheads, n ≥ 13 animals examined each). Scale bar represents 25 μm. e, Confocal sum projections of gonadal BM COL4A1/EMB-9::mRuby2, NID/NID-1::mNG, COL18A1/CLE-1::mNG, and PAPLN/MIG-6S::mNG in control and *unc-52* RNAi-treated 72-h adult animals (n = 5 animals examined each). Note the fibrillar structures (blue arrowheads) and aggregates (green arrowheads) of fluorescence within the gonadal BM upon knockdown of *unc-52*. Magenta dashed lines demarcate body wall muscle tissue where EMB-9::mRuby2 is produced. Scale bar represents 25 μm.

**Supplementary Figure 6.** TGFβ regulates BM collagen IV levels in *C. elegans*, and depletion of ADAMTS3 and ROBO impairs development and kidney BM function in zebrafish. A, Quantification of LAMC/LAM-2::mNG and COL4A1/EMB-9::mRuby2 gonadal BM fluorescence intensity upon knockdown of TGFβ ligand genes in *C. elegans* (n ≥ 20 each). b, Confocal z-slices of LAMC/LAM-2::mNG and COL4A1/EMB-9::mRuby2 (top) and quantification of fluorescence intensity (bottom) upon knockdown of TGF β type I receptor genes (n ≥ 20 each). Scale bar: 25 μm. c, Confocal z-slice of TGFβ type II receptor DAF-4::GFP depicting gonadal BM zone localization (outer tissue boundary in magenta). Scale bar represents 25 μm. d, Brightfield images of 2-days post fertilization (dpf) *col4a1*^gRNA^-injected (*col4a1* crispant) zebrafish embryos with intracerebral haemorrhage (asterisks). Scale bar represents 600 μm. e, Confocal images of collagen IV immunofluorescence (left) and brightfield images (right) of tail regions (dashed lines); arrows and arrowheads indicate reduced fin length and fin fold extension respectively in *robo1* and *adamts3* crispants (n = 5 animals examined each). f, Assessment of proteinuria (NL-D3 levels) in *robo1* and *adamts3* crispants (n = 24 each). For all boxplots, \*\*\*\**p*-value < 0.0001,\*\*\**p*-value < 0.001, \*\**p*-value < 0.01, ns—not significant; one-way ANOVA with post-hoc Dunnett’s test; edges indicate the 25^th^ and 75^th^ percentiles, the line in the box represents the median, and whiskers mark the minimum and maximum values.

**Supplementary Figure 7. Landscape of predicted loss-of-function variants in BM zone genes. a**, Classification of homozygous pLoF variants identified in 41 BM zone genes according to type of rare disease presented in associated 100KGP individuals. Count indicates number of variants observed in the respective genes. **b**, Odds ratio (OR) plots (comparing individuals with disease to unaffected relatives) for heterozygous and homozygous pre- dicted loss-of-function (pLoF) variants in BM zone genes grouped by their matrisome classification. \*\*\*\**p*-value < 0.0001, Fisher’s Exact test. **#**ORs for these homozygous pLoF variants could not be computed as there were no matched controls in 100KGP.

**Supplementary Figure 8.** Disease phenotypes observed in 100KGP individuals carrying *LAMA5*, *MATN2*, and *MPZL2* pLoF variants and functional analyses of *MATN2* variants. a, Disease phenotype associations for *LAMA5*, *MPZL2* and *MATN2* pLoF variants. Phenotypes observed in 100KGP individuals as well as previously reported phenotypes are shown. b, Western blots of MATN2 in lysate and ECM fractions derived from endogenous MATN2-depleted human podocytes over-expressing V5-tagged wild-type or variant MATN2. c, Top, human MATN2 protein domain structure. Bottom, 3D model of MATN2 EGF1 domain indicating disruption of the highlighted disulfide bond in the *MATN2^c.746G>C, p.Cys249Ser^* missense variant d, PCR amplification of the V5 tag using DNA extracted from podocytes transfected with the listed MATN2 constructs. PCR products amplified from the respective construct plasmids are shown as positive controls. No product was amplified from uninfected cells (negative control). e, *In vitro* splicing assay with a minigene for MATN2 variant 1 indicating aberrant splicing and the introduction of a premature stop codon (PTC).

**Supplementary table 1.** Protein databases used for the expansion of the BM zone network and verification of candidate BM localization.

**Supplementary table 2.** Conserved *C. elegans* matrisome genes with the BM gene expression signature.

**Supplementary table 3.** Interpro protein domains enriched in BM zone protein candidates.

**Supplementary table 4.** Candidate BM zone proteins identified by BM protein domain enrichment analysis and from predicted interaction with BM proteins.

**Supplementary table 5.** BM zone localization data for *C. elegans* orthologs of BM and CSI candidates fluorescently tagged in this and previous studies.

**Supplementary table 6.** Expanded list of BM zone gene network candidates.

**Supplementary table 7.** Verified BM zone gene network.

**Supplementary table 8.** BM zone protein abundance derived from human proteomic datasets.

**Supplementary table 9.** Human and mouse gene expression datasets used in this study.

**Supplementary table 10.** Expression variance of BM zone genes in human and mouse transcriptomic datasets.

**Supplementary table 11.** Connectivity metrics for human and *C. elegans* BM domain-based networks.

**Supplementary table 12.** Post-embryonic RNAi screen of 77 *C. elegans* BM zone gene orthologs.

**Supplementary table 13.** Human disease phenotype associations for BM zone genes.

**Supplementary table 14.** Top-level HPO terms associated with BM zone genes.

**Supplementary table 15.** BM zone gene variants identified in the 100KGP rare disease cohort.

**Supplementary table 16**. BM zone genes with homozygous pLoF variants in 100KGP cohort.

**Supplementary table 17**. Proteomic analysis of core BM protein levels in podocyte-derived matrix upon MATN2 knockdown.

**Supplementary table 18.** *C. elegans* and zebrafish strains used in this study.

**Supplementary table 19.** gRNA sequences used for CRISPR-Cas9-mediated knockdown in zebrafish.

**Supplementary table 20.** Primer sequences for minigene and PCR assays.

## Supplementary Notes

### Identification of basement membrane and cell surface interactor genes

To build a comprehensive network of basement membrane (BM) zone genes encoding BM proteins and cell surface interactors (CSI), we adopted the following sequential strategy: (i) initial identification of BM zone candidates through gene ontology; (ii) expansion of the BM gene zone network using five independent approaches; and (iii) verification of BM zone localization for candidates (detailed below and illustrated in **Fig. 1a**).

### Gene ontology retrieval

Gene Ontology (GO) Resource (release 2020-07)^30, 31^ was used to search for human genes catalogued under the ‘Basement membrane’ cellular component term (#GO:0008003/GO:0005605). A list of 103 human genes was obtained and used as the initial set of candidate BM zone genes for further analyses in this study. Within the 103 genes, 84 were present in the human matrisome^88^ gene list, including 64 structural genes, 17 matrix-associated components, and 3 BM receptors.

### Network expansion strategy

To expand the GO BM zone list, we established a curating strategy combining five approaches to capture other BM zone candidates not yet included in the human GO list, and also strategies to predict potential new BM zone proteins. The approaches included:

1) *Data curation*: We performed a systematic review of the literature and protein databases (listed in **Supplementary table 1**) to search for other putative BM components and cell surface interactors in vertebrates (human, rodents, zebrafish) and invertebrates (*C. elegans*, *Drosophila*) not included in the initial set of BM zone candidates. We used GO resource to screen for additional BM genes in mouse, and then identified and added the respective human orthologs. FASTA sequences for proteins encoded by candidate genes were obtained from UniProtKB and verified for (a) prediction of a signal peptide with PredSi (http://www.predisi.de/index.html)89 (b) transmembrane domains with the online tools TMHMM server v. 2.0 (http://www.cbs.dtu.dk/services/TMHMM/) and Phobius (https://phobius.sbc.su.se); and (c) evidence of secretion to the extracellular space and/or shedding from the cell surface for transmembrane proteins.

2) *BM gene expression signature in C. elegans*: Most BM protein encoding genes in *C. elegans* are synthesized in distant tissues, secreted into the extracellular fluid, and recruited to BMs for incorporation^90^. In particular, we noticed that 15 of the 18 known BM components in the worm (all BM genes listed previously^28^ except for *agr-1*, *pxn-2*, and *lam-1*) are synthesized in either the body wall muscles, or the body wall muscles and the epidermis (data derived from https://wormbase.org). We hypothesized that genes within the *C. elegans* matrisome^21^, that possess this expression signature could be BM gene candidates. Of 467 conserved matrisome genes, we identified 81 with the BM gene expression signature (**Supplementary table 2**).

3) *Conservation and enrichment analysis of BM protein domains*: We obtained a list of annotated putative BM protein domains for the genes in the expanding BM zone list from Interpro (https://www.ebi.ac.uk/interpro/; see **Supplementary table 3**). Using R software (R Core Team, 2020, https://www.R-project.org/), and the enricher function of the ClusterProfiler package (v3.14.3)^91^, we performed a domain enrichment analysis, taking the domain annotations for the entire human proteome as background. We identified 20 domains that were significantly over-represented in the expanding BM zone genes (adjusted *p*-value < 0.05, **Supplementary table 3**) and applied these domains as a filter onto the whole human proteome. We identified 25 additional BM zone gene candidates based on BM domain conservation (**Supplementary table 4**). Moreover, to determine whether human protein domains enriched in our BM zone list are conserved, we performed similar analyses for mouse, zebrafish, *Drosophila* and *C. elegans* ortholog genes.

4) *Nearest interacting neighbors of BM proteins*: To identify new BM zone gene candidates based on protein interaction, we searched for interacting partners of the proteins in the expanding BM zone list using the STRING (https://string-db.org/, version 11.0)^92^ database at a combined score threshold ≥ 70% (indicating high confidence interactions). This analysis returned 1533 BM interacting partners that were further filtered for presence of a signal peptide, enriched BM protein domains found in the previous analysis, and for direct interactions with at least 20 other BM proteins (**Supplementary table 4**). False positives (e.g., COL1A1, COL1A2, COL3A1) were manually removed and this analysis added 16 novel BM zone candidate genes.

5) *BM zone localization in C. elegans*: 36 genes have previously been localized to the BM zone in *C. elegans* (**Supplementary Table 5**, see genes tagged in previous studies), and human orthologs of these genes that were not already curated from the above analyses were added as candidates.

Together, these network expansion approaches identified 160 additional genes, resulting in a combined BM zone gene network comprising 263 candidates (**Supplementary table 6**).

### Verification of BM zone localization

To verify BM zone localization for the 263 genes in the expanded network, we first reviewed the literature and protein databases for evidence of tissue protein immunolocalization to BM zone in human, rodents, zebrafish, *Drosophila,* and *C. elegans*. Specifically, we used article search tools (NCBI PubMed, https://pubmed.ncbi.nlm.nih.gov/; Google Scholar, https://scholar.google.com/), the Human Protein Atlas (https://www.proteinatlas.org), and the Matrixome Project (http://dbarchive.biosciencedbc.jp/archive/matrixome/bm/home.html). Next, we fluorescently tagged a subset of *C. elegans* orthologs to investigate their localization, focusing on small gene families with clear orthology to human genes where none or only some of the family members have been localized to the BM zone in previous studies. We generated endogenous mNeonGreen or mRuby2 tags for 17 BM and 8 CSI candidate genes (**Methods** and **Supplementary Fig. 2**) and localized 14 of these genes to the BM zone (**Supplementary table 5**). Together, our literature review strategy and fluorescent tagging studies confirmed BM zone localization of 188 genes in the network (**Supplementary Fig. 1a**).

Among the remaining candidates without direct BM zone immunolocalization evidence, 34 were predicted to be in BM zone based on two metrics: high confidence interaction with one or more BM zone candidates (interaction data obtained from STRING at a combined score threshold ≥ 70%); and/or evidence for proteolytic activity on BM substrates (data collected from BRENDA, https://www.brenda-enzymes.org/; MEROPS, https://www.ebi.ac.uk/merops/; and publications; see **Supplementary Fig. 1a**).

We could not confirm or predict BM zone localization for 40 candidates due to insufficient evidence and/or GO mis-annotations (e.g., ALB, ANXA2, see **Figure 1a**). Thus, the integrated verified BM zone network comprised 163 BM components and 62 CSI. To validate this finalized BM zone list, we repeated the analysis for conservation and enrichment of BM protein domains (as described earlier). The validated list (**Supplementary table 7**) was used for subsequent analyses in this study. However, three components that were in later analyses were modified during the final proofing of the BM zone gene network prior to publication—PHF13 was a mis-annotation and was removed from the network; and HAPLN2 and MEGF6, which were erroneously classified under confirmed localization evidence, were transferred to the insufficient evidence category.

We adopted a colour code to distinguish components with confirmed (green) or predicted (magenta) localization to BM zone (**Fig. 1a**). Components in the insufficient evidence category were coloured in grey. Finally, to curate this network, we created an open access, interactive resource: ***b****asement **m**embrane*BASE (https://bmbase.manchester.ac.uk/).

### Identification of vertebrate and invertebrate orthologs

Orthologs for the 222 human BM zone genes were identified in mouse (*Mus musculus*) using MouseMine (http://www.mousemine.org/mousemine/begin.do), zebrafish (*Danio rerio*) using ZFIN (http://zfin.org/), fruit fly (*Drosophila melanogaster*) using FlyBase (<u>flybase.org</u>), and worm (*Caenorhabditis elegans*) using OrthoList 2 (http://ortholist.shaye-lab.org/) and WormBase (wormbase.org). We also used Ensembl (http://ensembl.org/index.html), NCBI (https://www.ncbi.nlm.nih.gov), and DIOPT (https://www.flyrnai.org/cgi-bin/DRSC_orthologs.pl) databases for verification. All orthologs are listed in **Supplementary table 7**.

### Case histories

***LAMA5***: *LAMA5* compound heterozygous c.3282+5G>A, and c.9489C>T;p.Tyr3163*. Chromosome array comparative genomic hybridization was normal for both fetuses, 7- dehydrocholesterol was normal in the first fetus. The first fetus was noted to have spina bifida, hypoplastic cerebellum, absent corpus callosum, ventricular septal defect, enlarged multicystic solitary left kidney, absent right kidney and urinary bladder, absent thymus, bilateral cleft lip and palate, laryngeal atresia, small ears with absent canals, cutaneous syndactyly 3-4th fingers and 2-3rd toes, exomphalos, imperforate anus and bilateral talipes. The second fetus was noted to have spina bifida, ventricular septal defect, bilateral renal agenesis, absent thymus, laryngeal stenosis, cutaneous syndactyly 2-3rd and partial syndactyly 3-4th fingers, syndactyly 2-3-4th toes and anteriorly placed anus. Negative panel screening for CAKUT v1.4 and specifically Fraser syndrome, and also screened for arthrogryposis v3, clefting v2.0 disorder of sex development v2.1, hydrocephalus v2.1, intellectual disability v3.0, limb disorder v2.0 malformations of cortical development v2.2, non-syndromic familial congenital anorectal malformations v1.6, and VACTERL-like phenotype v1.25. Both pregnancies were terminated.

***MPZL2***: The case (homozygous c.72del) was father of the individual (daughter and proband) recruited to the 100KGP. The case was diagnosed with cystic kidney disease at the age of 35. The daughter (c.72del heterozyote) had multiple renal cysts diagnosed at the age of 15 and no other phenotypes described. It was not possible to obtain further clinical information from the recruiting clinician.

***MATN2***: Case 1 (C.1081+3_1081+6del): male proband with early onset or familial intestinal pseudo-obstruction at the age of >1 year; Case 2 (c.1585del): male proband recruited at the age of >1 year into Ultra-rare disorder Disease Group (from the Genomics England 100,000 genomes project); Case 3 (c.746G>C and C.1450+1G>A case 3): male proband diagnosed with intellectual disability at the age of 6. It was not possible to obtain further clinical information from the recruiting clinician.

## Notes

https://doi.org/10.6084/m9.figshare.c.5662348

### Competing Interest Statement

The authors have declared no competing interest.

