## Supplementary fig. 1 for "A basement membrane discovery pipeline uncovers network complexity, new regulators, and human disease associations"

Supplementary Figure 1

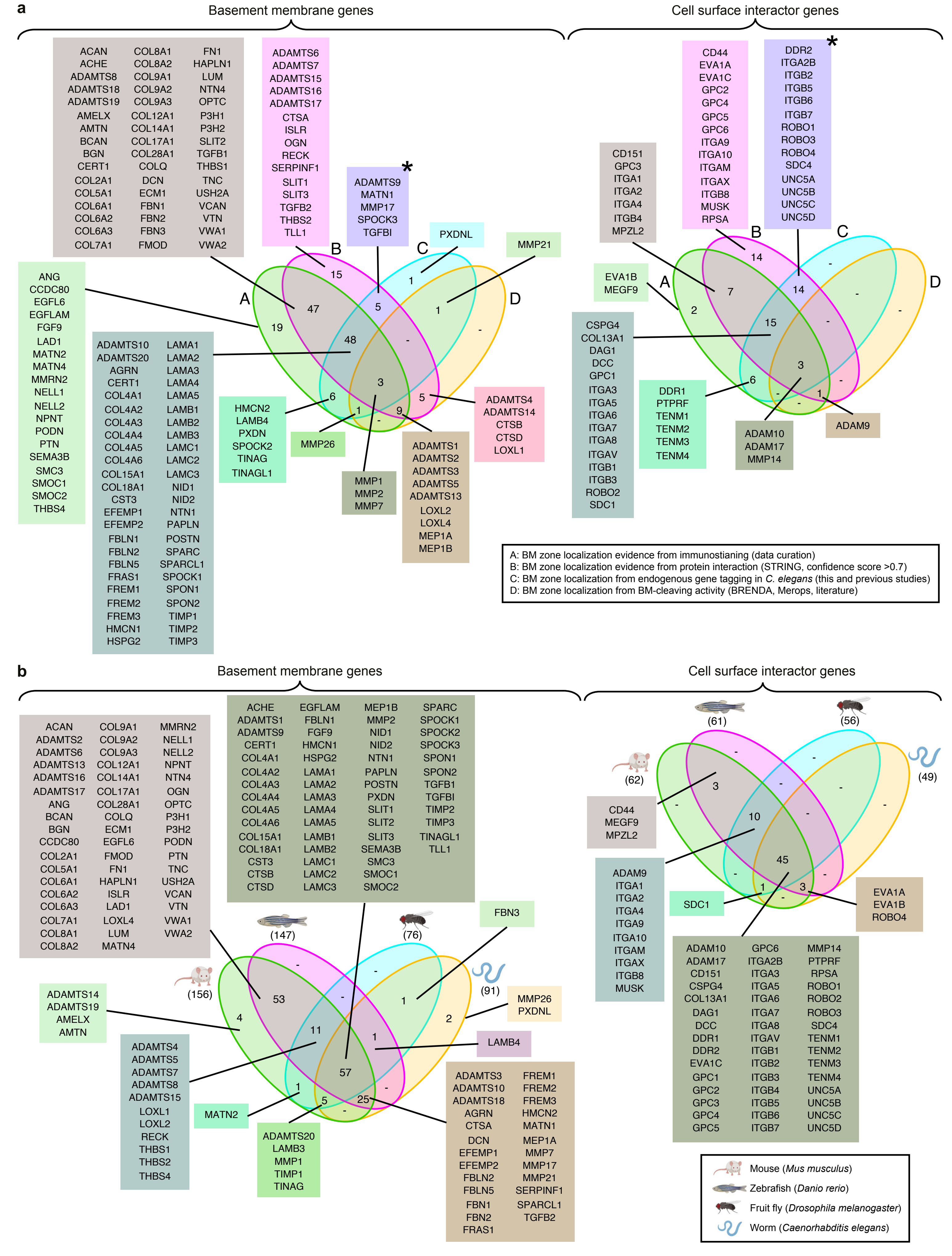


**Supplementary Figure 1. Verification strategies for basement membrane zone localization and conservation of network genes. a**, Venn diagrams illustrate how the localization of human proteins in the integrated basement membrane (BM) zone network was verified. The asterisk indicates a group of proteins with predicted localization whose *C. elegans* orthologs were detected in the BM zone through fluorescent tagging. **b**, Human BM matrix and cell surface interactor (CSI) genes are shown in boxes and the presence of corresponding orthologs in mouse, zebrafish, *D. melanogaster*, and *C. elegans* (Supplementary Fig. 3 and Supplementary table 7) is represented in Venn diagrams. Animal illustrations made with https://biorender.com.
