## Supplementary fig. 2 for "A basement membrane discovery pipeline uncovers network complexity, new regulators, and human disease associations"

Supplementary Figure 2

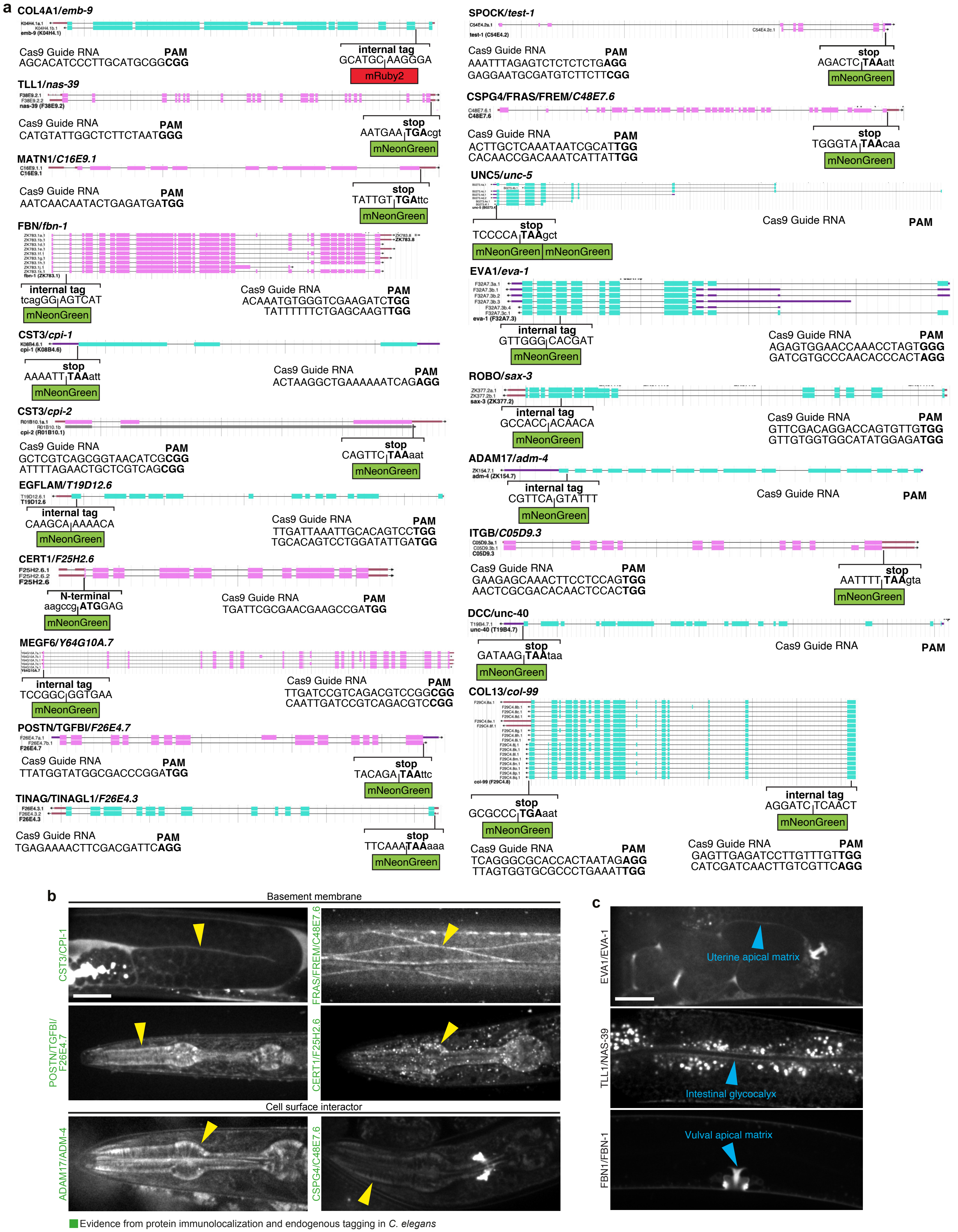


**Supplementary Figure 2. Visualization of basement membrane zone candidate localization in *C. elegans* through fluorescent protein insertions at endogenous gene loci.** **a**, Worm gene schematics (adapted from https://wormbase.org) indicate genomic locations where mNeonGreen (green box), two tandem mNeonGreen (two green boxes), or mRuby2 (red box) fluorophores were inserted for BM and CSI candidates tagged in this study. For each gene, sgRNA, PAM site, and insertion site sequences are provided. Every gene was tagged at a single locus, except for *col-99*, which was tagged at two separate loci. Note that exons are colored magenta or cyan depending on alignment to the forward or reverse strand of the reference genome, respectively. **b**, Confocal middle-plane z-slices of tagged candidates in adult animals showing BM zone localization (yellow arrowheads) to either the pharyngeal or gonadal BM, except for FRAS/FREM/C48E7.6, which localizes to the body wall muscle BM. **c**, Confocal z-slices of candidates not detected in the BM zone but present in other matrices (blue arrowheads). Scale bar represents 25 μm.
