## Supplementary fig. 3 for "A basement membrane discovery pipeline uncovers network complexity, new regulators, and human disease associations"

Supplementary Figure 3

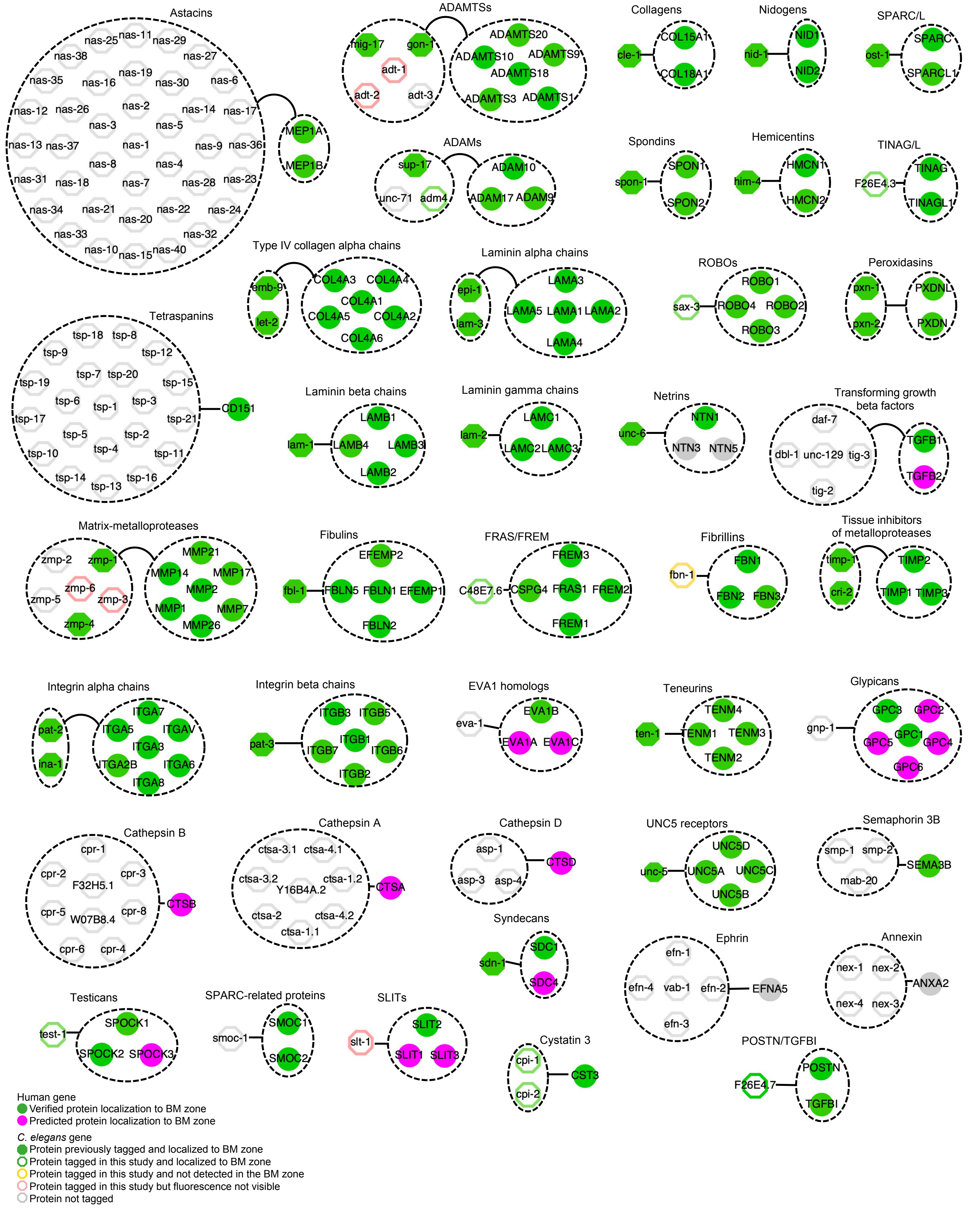


**Supplementary Figure 3. Basement membrane zone gene families in *C. elegans*.** Diagrams illustrate the orthologous relationships of *C. elegans* genes and gene families (on the left) with human BM zone genes (on the right). Note the various one-to-many and many-to-many relationships between several genes for both species.
