## Supplementary fig. 4 for "A basement membrane discovery pipeline uncovers network complexity, new regulators, and human disease associations"

Supplementary Figure 4

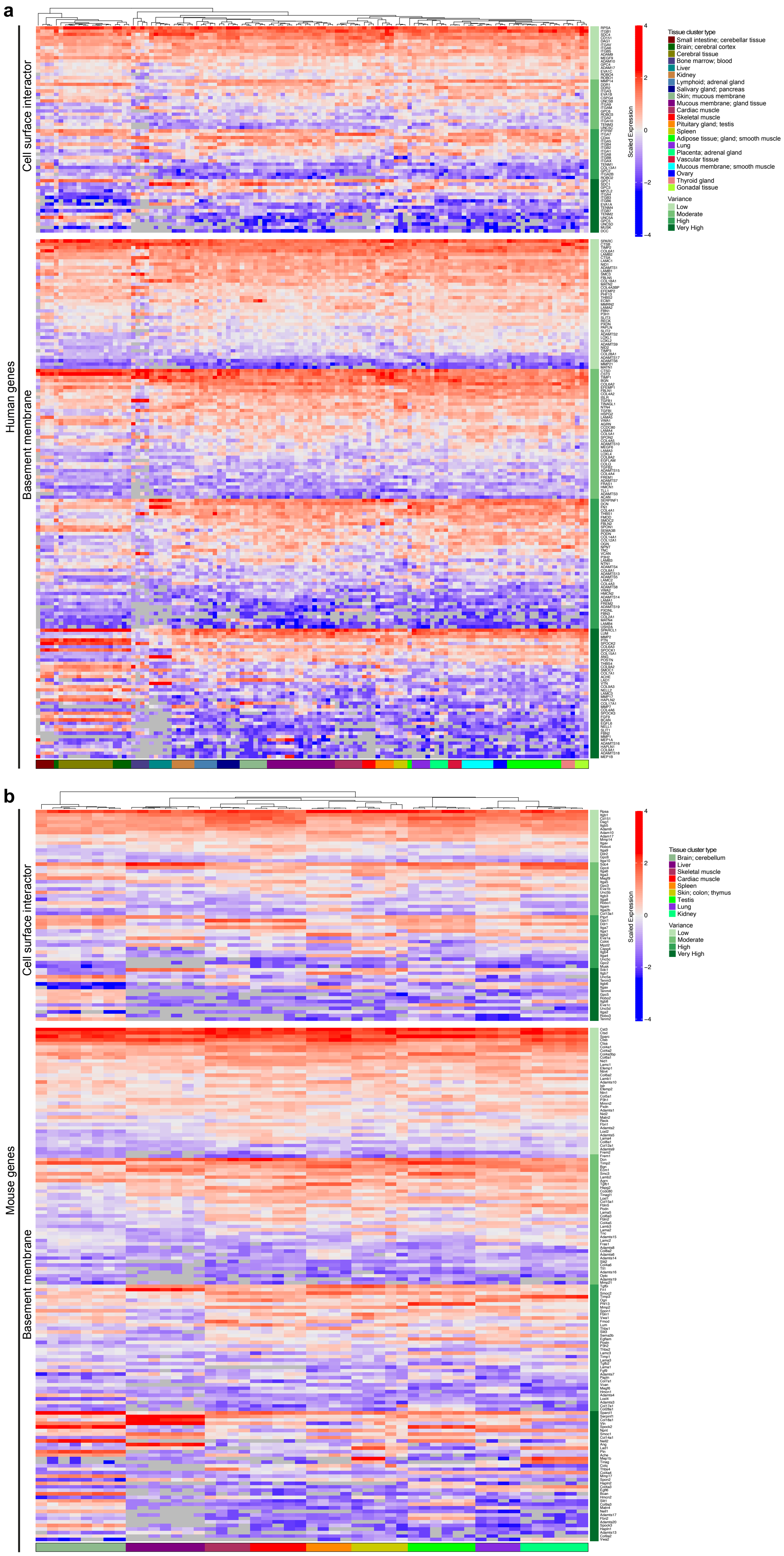


**Supplementary Figure 4. Expression variance of mammalian basement membrane zone genes.** Heatmaps indicate scaled gene expression of BM matrix and CSI genes across different (a) human and (b) mouse tissues (transcriptomic datasets used in this study detailed in Supplementary table 9). Genes are binned according to expression variance (low, moderate, high, very high; Supplementary table 10).
