## Supplementary fig. 5 for "A basement membrane discovery pipeline uncovers network complexity, new regulators, and human disease associations"

Supplementary Figure 5

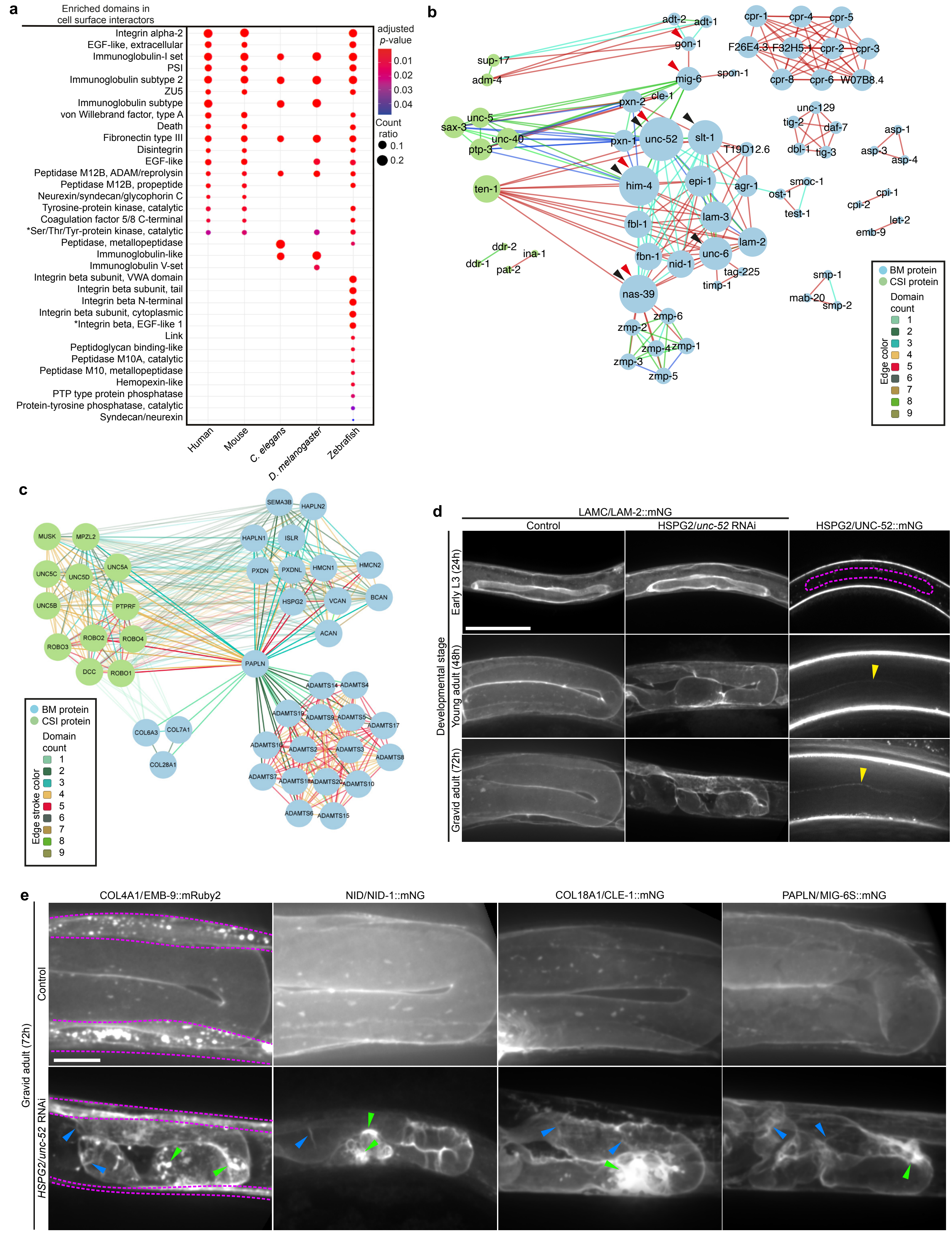


**Supplementary Figure 5. Perlecan/UNC-52 and papilin/MIG-6 are hub proteins and depletion of *C. elegans* UNC-52 disrupts basement membrane organization. a,** Conservation and enrichment of Interpro protein domains (as described in Fig. 3a) in verified CSI components. **b**, A domain-based interactome (as described in Fig. 3b) for *C. elegans* BM zone genes. **c**, Sub-network highlighting diverse groups of human BM zone proteins that share domains with papilin. **d**, Left, confocal sum projections of gonadal BM LAMC/LAM-2::mNG in control and *unc-52* RNAi-treated early L3, young adult, or gravid adult animals (n = 10 animals examined each). Right, confocal middle-plane z-slices of HSPG2/UNC-52::mNG, which is not detected at the gonadal BM (outlined in magenta) in the early L3 stage, but is visible in the young adult and gravid adult stages (yellow arrowheads, n ≥ 13 animals examined each). Scale bar represents 25 μm. **e**, Confocal sum projections of gonadal BM COL4A1/EMB-9::mRuby2, NID/NID-1::mNG, COL18A1/CLE-1::mNG, and PAPLN/MIG-6S::mNG in control and *unc-52* RNAi-treated 72-h adult animals (n = 5 animals examined each). Note the fibrillar structures (blue arrowheads) and aggregates (green arrowheads) of fluorescence within the gonadal BM upon knockdown of *unc-52*. Magenta dashed lines demarcate body wall muscle tissue where EMB-9::mRuby2 is produced. Scale bar represents 25 μm.
