## Supplementary fig. 6 for "A basement membrane discovery pipeline uncovers network complexity, new regulators, and human disease associations"

Supplementary Figure 6

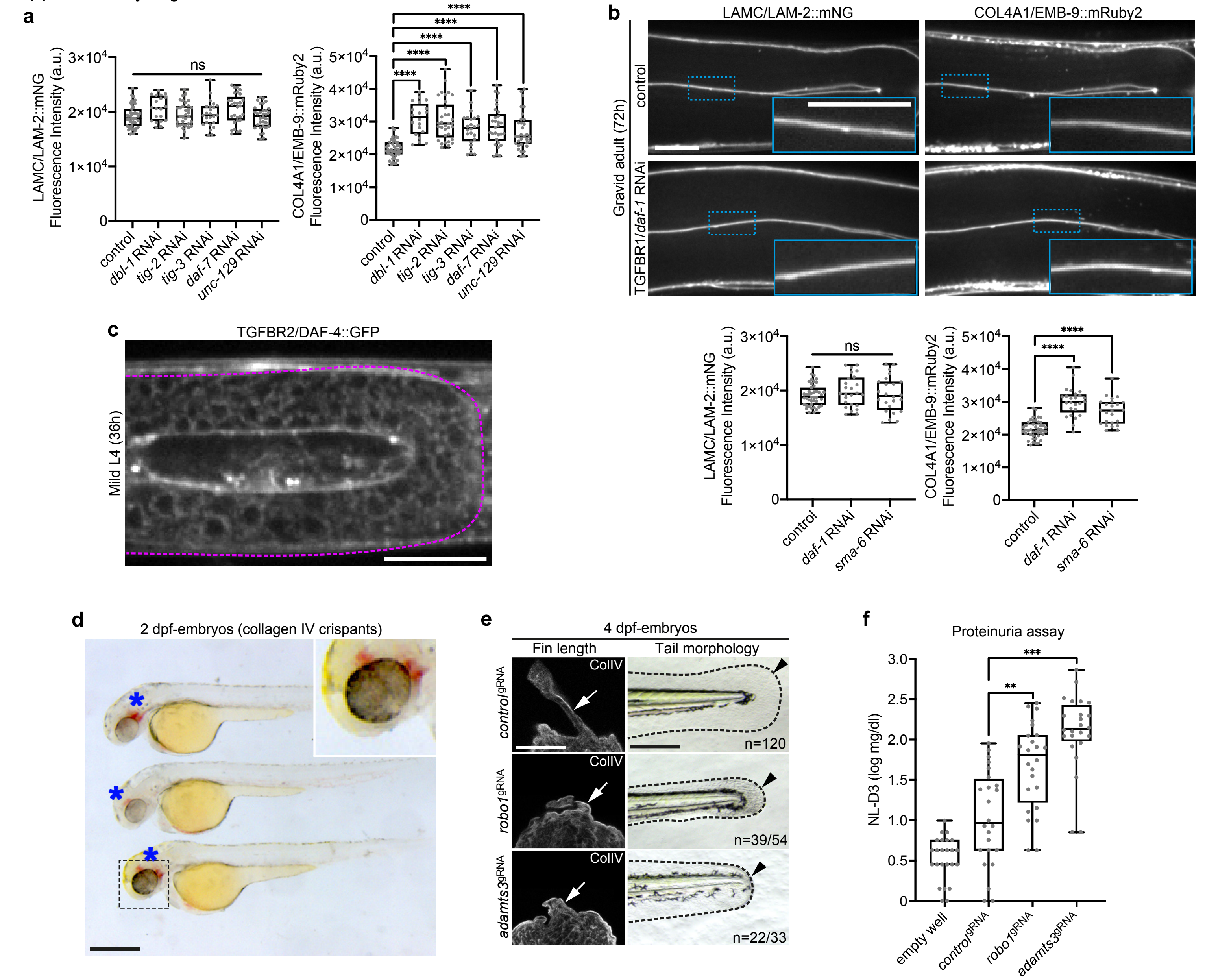


**Supplementary Figure 6. TGFβ regulates BM collagen IV levels in *C. elegans*, and depletion of ADAMTS3 and ROBO impairs development and kidney BM function in zebrafish. a,** Quantification of LAMC/LAM-2::mNG and COL4A1/EMB-9::mRuby2 gonadal BM fluorescence intensity upon knockdown of TGFβ ligand genes in *C. elegans* (n ≥ 20 each). **b**, Confocal z-slices of LAMC/LAM-2::mNG and COL4A1/EMB-9::mRuby2 (top) and quantification of fluorescence intensity (bottom) upon knockdown of TGFβ type I receptor genes (n ≥ 20 each). Scale bar: 25 µm. **c**, Confocal z-slice of TGFβ type II receptor DAF-4::GFP depicting gonadal BM zone localization (outer tissue boundary in magenta). Scale bar represents 25 µm. **d**, Brightfield images of 2-days post fertilization (dpf) *col4a1*$^{gRNA}$-injected (*col4a1* crispant) zebrafish embryos with intracerebral haemorrhage (asterisks). Scale bar represents 600 µm. **e**, Confocal images of collagen IV immunofluorescence (left) and brightfield images (right) of tail regions (dashed lines); arrows and arrowheads indicate reduced fin length and fin fold extension respectively in *robo1* and *adamts3* crispants (n = 5 animals examined each). **f**, Assessment of proteinuria (NL-D3 levels) in *robo*1 and *adamts3* crispants (n = 24 each). For all boxplots, *****p*-value < 0.0001,****p*-value < 0.001, ***p*-value < 0.01, ns—not significant; one-way ANOVA with post-hoc Dunnett's test; edges indicate the 25th and 75th percentiles, the line in the box represents the median, and whiskers mark the minimum and maximum values.
