## Supplementary fig. 7 for "A basement membrane discovery pipeline uncovers network complexity, new regulators, and human disease associations"

Supplementary Figure 7

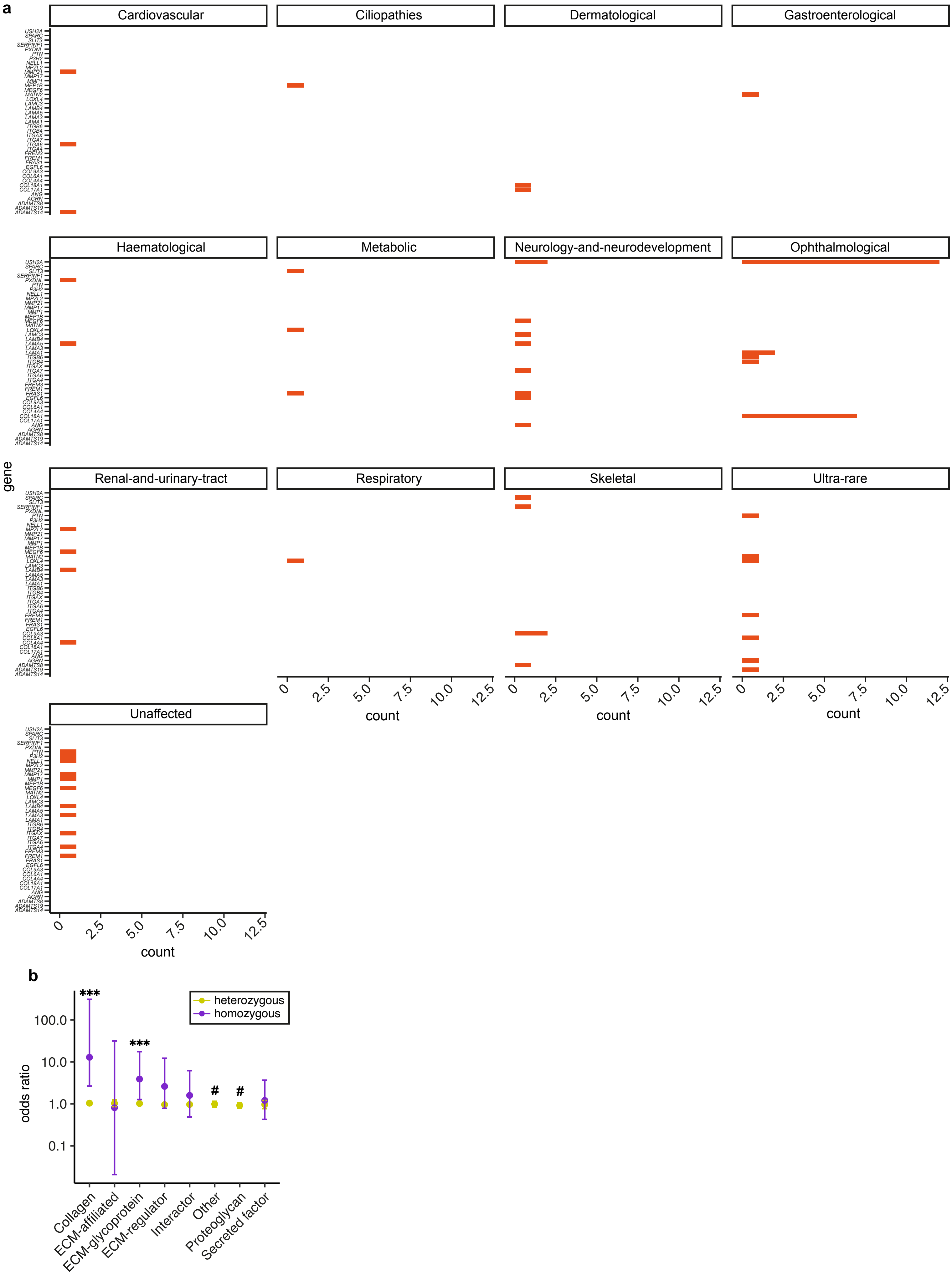


**Supplementary Figure 7. Landscape of predicted loss-of-function variants in BM zone genes. a**, Classification of homozygous pLoF variants identified in 41 BM zone genes according to type of rare disease presented in associated 100KGP individuals. Count indicates number of variants observed in the respective genes. **b**, Odds ratio (OR) plots (comparing individuals with disease to unaffected relatives) for heterozygous and homozygous predicted loss-of-function (pLoF) variants in BM zone genes grouped by their matrisome classification. ****$p$-value < 0.0001, Fisher's Exact test. [#]ORs for these homozygous pLoF variants could not be computed as there were no matched controls in 100KGP.
