## Supplementary fig. 8 for "A basement membrane discovery pipeline uncovers network complexity, new regulators, and human disease associations"

Supplementary Figure 8

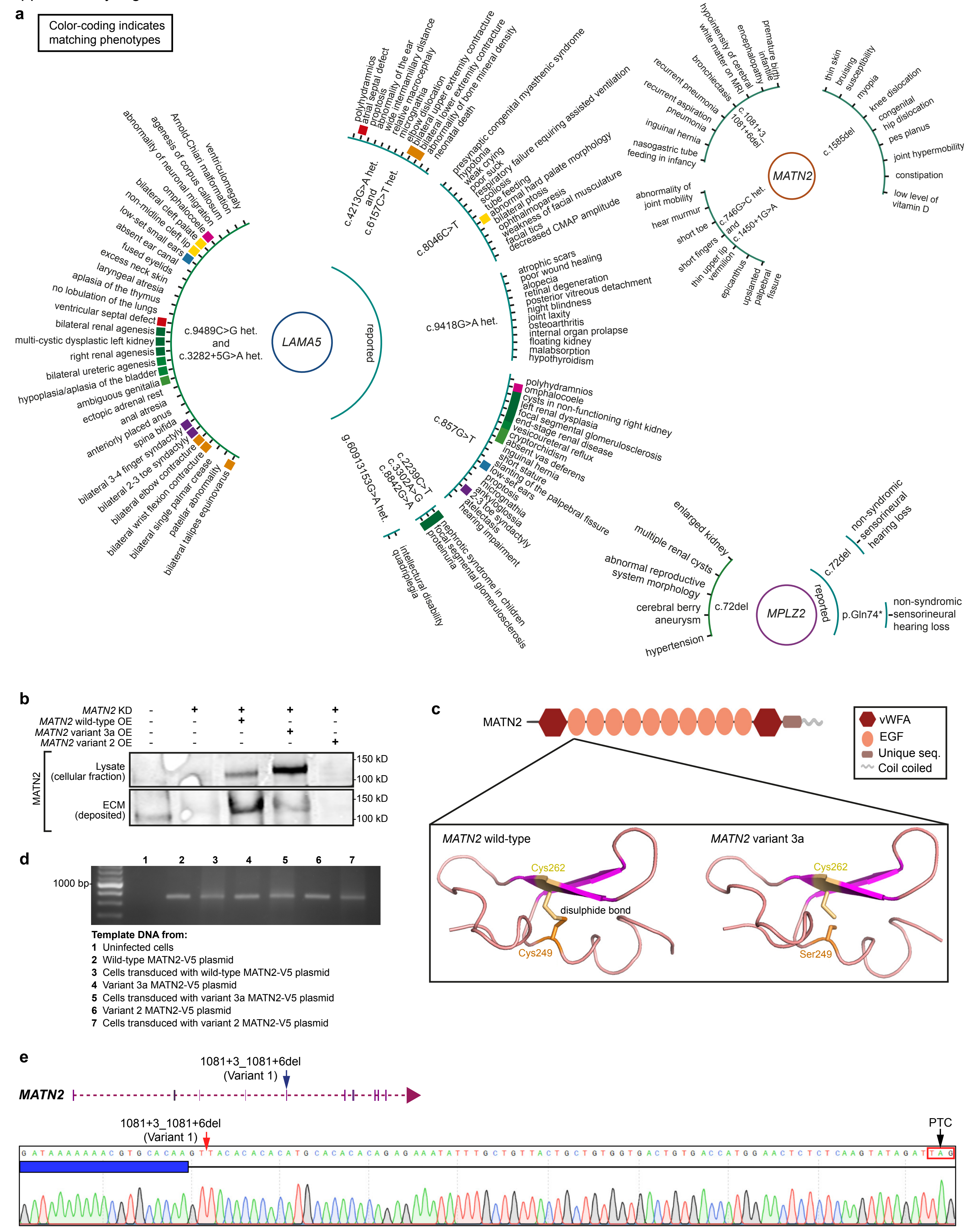

**Supplementary Figure 8. Disease phenotypes observed in 100KGP individuals carrying *LAMA5*, *MATN2*, and *MPZL2* pLoF variants and functional analyses of *MATN2* variants. a,** Disease phenotype associations for *LAMA5*, *MPZL2* and *MATN2* pLoF variants. Phenotypes observed in 100KGP individuals as well as previously reported phenotypes are shown. **b**, Western blots of MATN2 in lysate and ECM fractions derived from endogenous MATN2-depleted human podocytes over-expressing V5-tagged wild-type or variant *MATN2*. **c**, Top, human MATN2 protein domain structure. Bottom, 3D model of MATN2 EGF1 domain indicating disruption of the highlighted disulfide bond in the *MATN2*$^{c.746G>C,\ p.Cys249Ser}$ missense variant. **d**, PCR amplification of the V5 tag using DNA extracted from podocytes transduced with the listed MATN2 constructs. PCR products amplified from the respective construct plasmids are shown as positive controls. No product was amplified from uninfected cells (negative control). **e**, *In vitro* splicing assay with a minigene for *MATN2* variant 1 indicating aberrant splicing and the introduction of a premature stop codon (PTC).
